# IL-2 and CD40L can establish and sustain a human B cell-mediated immune regulatory network

**DOI:** 10.64898/2026.10.12.755729

**Authors:** Laurie Baert, Adrien Caffiers, Agnes Yang, Sae Rome Choi, Suman Bose, SangKon Oh

## Abstract

B cells can control inflammation, but the factors governing B cell-mediated immune regulatory network remain unclear. We report that IL-2/CD40L enable class-switched memory (CSM) B cells to acquire an immunosuppressive phenotype characterized by the expression of surface TIGIT as well as multiple other inhibitory receptors (PD-1, LAG-3, GITR, CD39, and CTLA-4) and soluble molecules (IL-10, granzyme B, and IL-35). TIGIT^+^ CSM B cells expressing functional CCR9 exhibit significant defects in total immunoglobulin secretion, while preserving IgA2 secretion. Furthermore, IL-2/CD40L-induced *ex vivo*-generated TIGIT^+^ CSM B cells suppress human T cell-mediated inflammation in a xenogeneic mouse model of acute graft-versus-host disease, while promoting IL-10-producing regulatory T cell (Treg) response. These findings identify IL-2/CD40L as key drivers of a B cell-mediated immune regulatory network, especially by generating TIGIT^+^ CSM B cells that can further promote Treg response. This study supports the development of TIGIT^+^ B cell-based therapy models for inflammatory diseases.

**Graphical Abstract:** 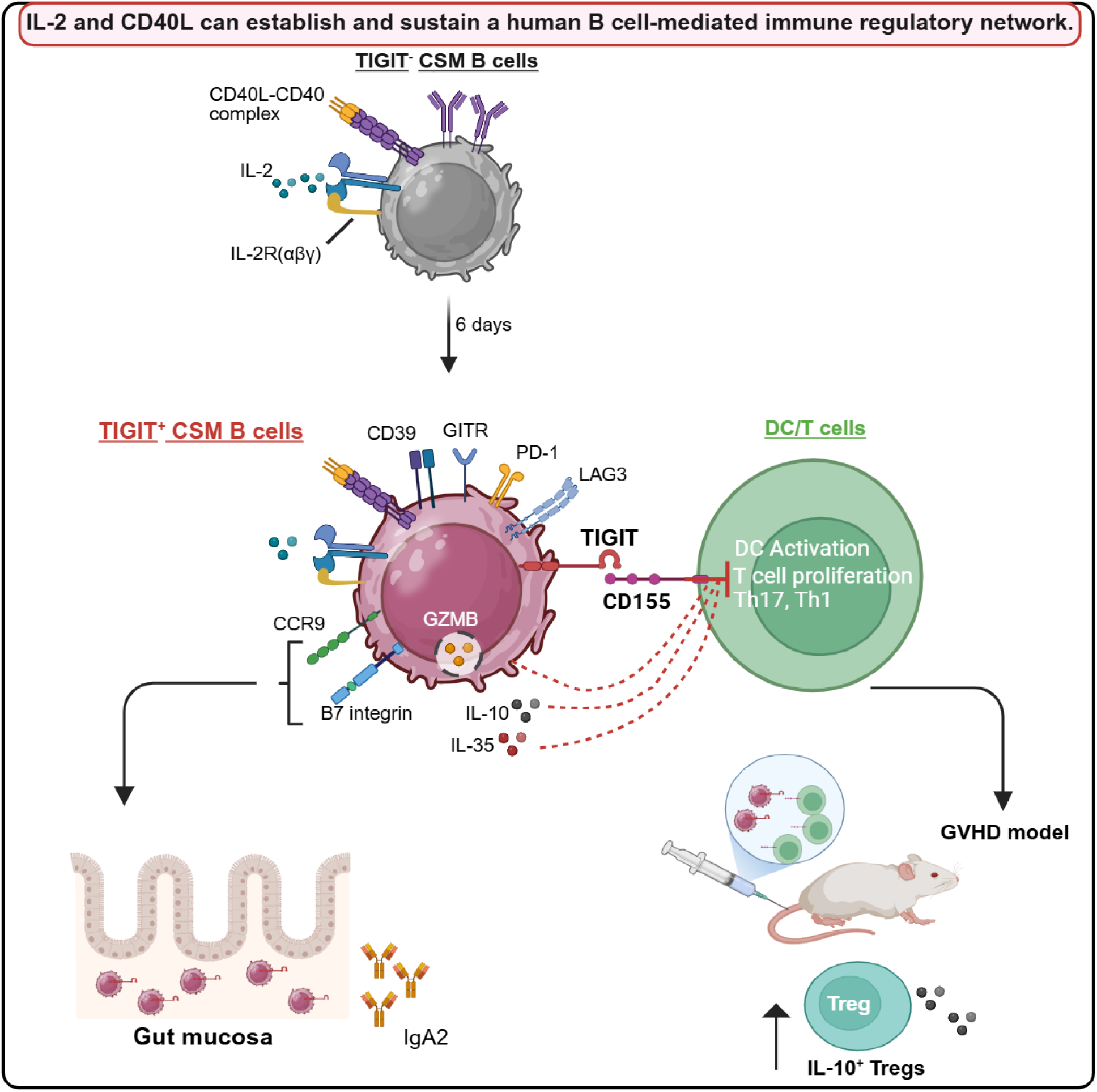

**Highlights:**

- IL-2/CD40L can generate and maintain immunosuppressive TIGIT^+^ B cells in humans.
- TIGIT^+^ B cells expressing CCR9 have defects in antibody secretion, but not IgA2.
- TIGIT^+^ B cells suppress inflammation, while promoting Treg response *in vivo*.
- IL-2/CD40L can thus build a robust B cell-mediated immune regulatory network.

## INTRODUCTION

B cell-mediated immune regulation has been reported in autoimmune disease^1–6^, transplantation rejection,^7–11^ microbial infection^12–16^ and cancer.^5,17–20^ Nonetheless, how B cells can acquire immune regulatory functions remains poorly understood. Unlike regulatory T cells (Tregs), there is no lineage specific marker representing immunosuppressive B cells. IL-10 expression has thus been used as a hallmark of immunosuppressive B cells. However, different subsets of B cells can control inflammatory responses by expressing not only IL-10, but also other inhibitory molecules, including IL-35 and granzyme B, as well as cell surface molecules, such as PD-L1, PD-1, CD39, CD73, and CTLA-4.^12,21–26^

We recently reported that a small fraction of B cells in the blood of healthy individuals express a co-inhibitory receptor, T cell immunoreceptor with Ig and ITIM domains (TIGIT).^27^ TIGIT^+^ B cells can control inflammatory T cell responses, including Th1, Th17, and T follicular helper (Tfh) cell responses, not only by directly interacting with T cells, but also by controlling inflammatory function of dendritic cells (DCs),^9,27^ which has a high potential to result in substantial consequences. Indeed, an impaired TIGIT expression on B cells leads to the increases of blood circulating Tfh cells in multiple sclerosis (MS) patients,^6^ contributing to the pathogenesis of MS. Although subsets of B cells expressing TIGIT are not the same in mice and humans, e.g., TIM-1^+^ B cells in mice^28^ versus memory B cells^6^ especially class-switched memory (CSM) B cells^27^ in humans, TIGIT deficiency in B cells also leads to spontaneous inflammation in central nervous system (CNS) in mice.^28^ Furthermore, the frequency of blood circulating TIGIT^+^ B cells correlated with the frequency of Tregs, while inversely correlating with the frequency of Tfh cells in renal allograft recipients.^9,27^ Notably, the outcomes of reduced or lack of TIGIT expression on B cells^6,27,28^ are observed despite the presence of TIGIT-expressing T, NK, and NKT cells,^29–33^ suggesting that TIGIT^+^ B cells might possess unique immune regulatory functions that cannot be compensated by other TIGIT-expressing cells.

To understand such novel immune regulatory function of TIGIT^+^ B cells, this study investigated key biological factors that could govern TIGIT expression on human B cells. We found that the CD4^+^ helper T cell-driven factors, IL-2 and CD40L, were the major contributors for the generation and maintenance of TIGIT^+^ CSM B cells. Notably, TIGIT^+^ CSM B cells also upregulate a variety of currently known B cell-derived inhibitory molecules (i.e., PD-1, CD39, LAG-3, GITR, and CTLA-4 as well as IL-10, IL-35, and granzyme B) while having significant defects in immunoglobulin (Ig) secretion. TIGIT^+^ CSM B cells further display unique molecular and cellular features, supporting their potent immune regulatory functions and their migration towards specific mucosal tissues. Finally, we validated the immune regulatory function of IL-2/CD40L-induced *ex vivo*-generated TIGIT^+^ CSM B cells using a xenogeneic mouse model of acute graft-versus-host disease (aGVHD). Mechanistic insight into the TIGIT^+^ CSM B cell-mediated immune regulation has also been investigated.

## RESULTS

### IL-2 and CD40L can induce TIGIT expression in CSM B cells

To identify factors associated with TIGIT expression in B cells, we analyzed RNA-seq data of CSM B cells and transitional B cells (TBs) purified from the blood of six healthy individuals.^27^ As expected, human *TIGIT* expression was highly enriched in CSM B cells compared with TBs (**Figure 1A**), consistent with previous reports demonstrating that TIGIT is primarily expressed by memory B cells, including CD24^hi^CD27^+^ B10-like B cells.^6,27^ In contrast, unlike murine TIM-1^+^ B cells,^34^ TIM-1^+^ TBs in adult human blood express little or no *TIGIT* (**Figure 1A**).^27^

**Figure 1.**
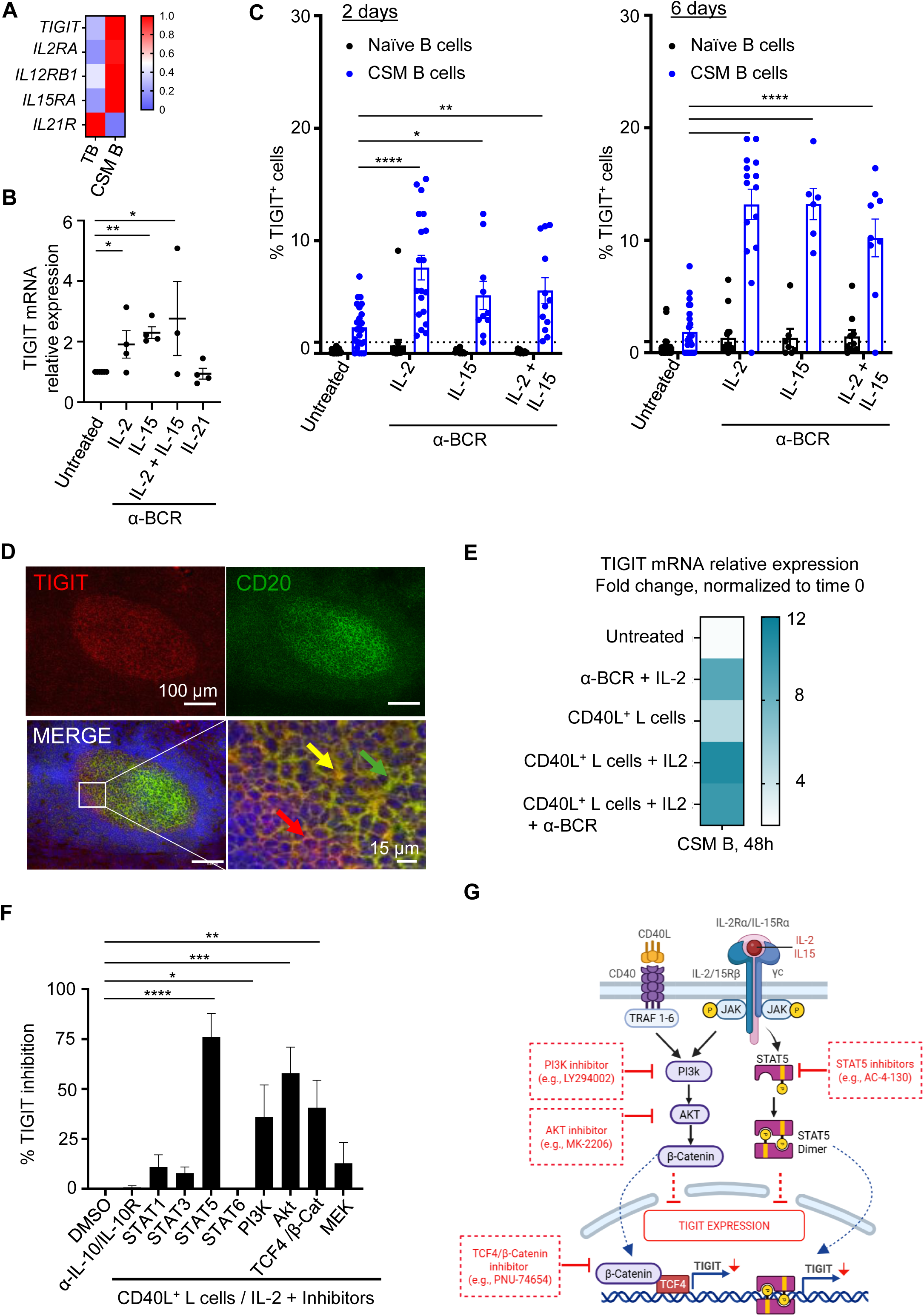
IL-2 and CD40L induce TIGIT expression in CSM B cells via the activation of STAT5 and PI3K/AKT pathways. **(A)** Heatmap showing the expression levels of *TIGIT, IL2RA*, *IL12RB1*, *IL15RA*, *and IL21R* from RNA-seq analysis of transitional B cells (TBs) and CSM B cells purified from the blood of healthy adult individuals (N = 6). **(B)** Anti-BCR-activated CD19^+^ total B cells were cultured 36 h in the presence of indicated cytokines. *TIGIT* mRNA expression levels were measured by qPCR assay. Each dot represents data generated with cells from individual donors (N = 3-4). **(C)** Anti-BCR-activated CD19^+^ B cells were cultured for 2 and 6 days in the indicated conditions. The frequency of TIGIT^+^ B cells in naïve (CD19^+^CD27^−^IgD^+^) and CSM B cells (CD19^+^CD27^+^IgD^−^) was measured by flow cytometry. **(D)** Fixed frozen tonsil sections were stained with DAPI (blue), anti-TIGIT (red) and anti-CD20 (green) antibodies (20x; Bar size 100 µm). The rectangle in the merged image indicates the magnified area (Bar size 15 µm). Images acquired with control antibodies are in Figure S4. **(E)** Purified CSM B cells from healthy donors (N = 4-8) were cultured with indicated stimuli for 48 h, and *TIGIT* mRNA expression levels were measured by qPCR assay. **(F)** Purified CSM B cells (N = 4-6) were pre-incubated with indicated inhibitors for 1-2 h and then cultured in the presence of IL-2 and γ-irradiated CD40L-L cells for 6 days. The percentage of TIGIT^+^ CSM B cells was measured by flow cytometry. Data are presented as the percentage of TIGIT inhibition compared to the control, DMSO alone. **(G)** Proposed pathways for the IL-2/CD40L-induced TIGIT expression in CSM B cells. Data are presented as mean ± SEM. Statistical analyses were performed using one-way ANOVA with Dunnett’s multiple comparisons test (**B** and **F**) and two-way ANOVA with Tukey’s multiple comparisons test (**C**). ∗ <0.05, ∗∗ <0.01, ∗∗∗ <0.001, ∗∗∗∗ <0.0001.

In addition to elevated *TIGIT* expressions, CSM B cells also upregulated expression of both *IL-2RA* and *IL-15RA* mRNA transcripts, suggesting their enhanced responsiveness to common γ-chain cytokines, IL-2 and IL-15, respectively. In opposite, CSM B cells, when compared to TBs, downregulated *IL-21R* expression. (**Figure 1A**).

We next tested whether those γ-chain cytokines could affect *TIGIT* mRNA expression in anti-BCR-activated B cells (**Figure 1B**). Both IL-2 and IL-15 enhanced *TIGIT* mRNA expression, but combination of IL-2 and IL-15 did not further increase *TIGIT* expression. Neither IL-21 (**Figure 1B**) nor IL-12p40 (not shown) affected *TIGIT* mRNA expression, although IL-21 increased *PRDM1* expression (**Figure S1A**). Unlike in T cells,^35–38^ IL-10 or IL-27 did not promote *TIGIT* mRNA expression in B cells, but IL-10 increased *PRDM1* expression (**Figure S1A**). Pam3CSK4 (TLR2 ligand),^39^ Poly (I:C) (TLR3 ligand),^40,41^ or CpG ODN2006 (TLR9 ligand)^42^ did not increase *TIGIT* mRNA expression in B cells (**Figure S1B**). Therefore, we concluded that IL-2 and IL-15 contribute to *TIGIT* mRNA expression in anti-BCR-activated B cells. Anti-BCR alone did not significantly affect *TIGIT* mRNA expression (**Figure S2**).

In line with *TIGIT* mRNA expression, both IL-2 and IL-15 increased surface TIGIT expression on anti-BCR-activated CSM B cells after 2 days and higher percentage was observed after 6 days of culture (**Figure 1C and Figure S3**). Neither IL-2 nor IL-15 significantly affected surface TIGIT expressions on naïve B cells, which is consistent with previous observations.^6,27^

As CD4^+^ helper T cells are the major sources of IL-2, we hypothesized that TIGIT^+^ B cells could be found in lymphoid tissues where B cells interact with CD4^+^ helper T cells. To test this hypothesis, inflamed human tonsil sections were stained with anti-TIGIT and anti-CD20 antibodies (**Figure 1D and Figure S4**). Indeed, CD20^+^ B cells expressing TIGIT were found mainly at the interface between B cell follicles and surrounding T cell zones, supporting the physiological relevance of TIGIT expression on B cells in the lymphoid tissues.

In consideration of the T-B cell interactions (**Figure 1D**), we next asked whether CD40L could also contribute to TIGIT expression by CSM B cells. As shown in **Figure 1E**, stimulation with CD40L modestly increased *TIGIT* mRNA expression compared with untreated cells. The addition of IL-2 markedly enhanced CD40L-induced TIGIT expression, resulting in the highest *TIGIT* mRNA levels, exceeding those induced by anti-BCR and IL-2. Anti-BCR stimulation did not further increase *TIGIT* mRNA expression beyond that induced by the combination of CD40L and IL-2.

The roles of IL-2 and CD40L in the induction of TIGIT expression on CSM B cells were further confirmed in **Figure 1F**. IL-2 can signal through STAT5,^43,44^ and STAT5 inhibitor significantly inhibited TIGIT expression on CSM B cells. Ligation of CD40 is known to activate PI3K-AKT signaling pathway that can lead to the activation of β-catenin/TCF4.^45^ **Figure 1F** shows that inhibitors of PI3K, AKT, and TCF4/β-catenin also decreased surface TIGIT expression on CSM B cells. Although IL-10 was reported to induce TIGIT expression in T cells,^35,36,38^ blocking IL-10 or MEK/ERK inhibitors did not affect TIGIT expression by CSM B cells, which is consistent with the data in **Figure S1A**.

Collectively, combination of IL-2 and CD40L can induce TIGIT expression in human CSM B cells via the activation of STAT5- and PI3K/AKT-mediated signaling pathways, as summarized in **Figure 1G**.

### CD40L helps maintain surface TIGIT expression on CSM B cells

To further examine the roles of IL-2, CD40L, and combination of IL-2 and CD40L in the generation and maintenance of TIGIT^+^ CSM B cells, we measured the frequency as well as actual numbers of TIGIT^+^ B cells after culturing CSM B cells for 6 days in different conditions.

In line with the data in **Figure 1E**, IL-2 increased the percentage of TIGIT^+^ CSM B cells, but this was not observed in the absence of anti-BCR stimulation (**Figure 2A**). We also found that CD40L significantly increases the percentage of TIGIT^+^ CSM B cells in all culture conditions. CD40L alone was also sufficient to significantly increase the percentage of TIGIT^+^ B cells on day 6 while the addition of anti-BCR did not further increase TIGIT expression (**Figure 2A and Figure S5**). Consequently, combination of CD40L with IL-2 and CD40L alone resulted in the highest absolute numbers of TIGIT^+^ CSM B cells on day 6 (**Figure 2B**), and this was explained by the data in **Figure 2C**, showing that TIGIT^+^ B cells proliferate better than TIGIT^−^ B cells in the presence of CD40L and IL-2.

**Figure 2.**
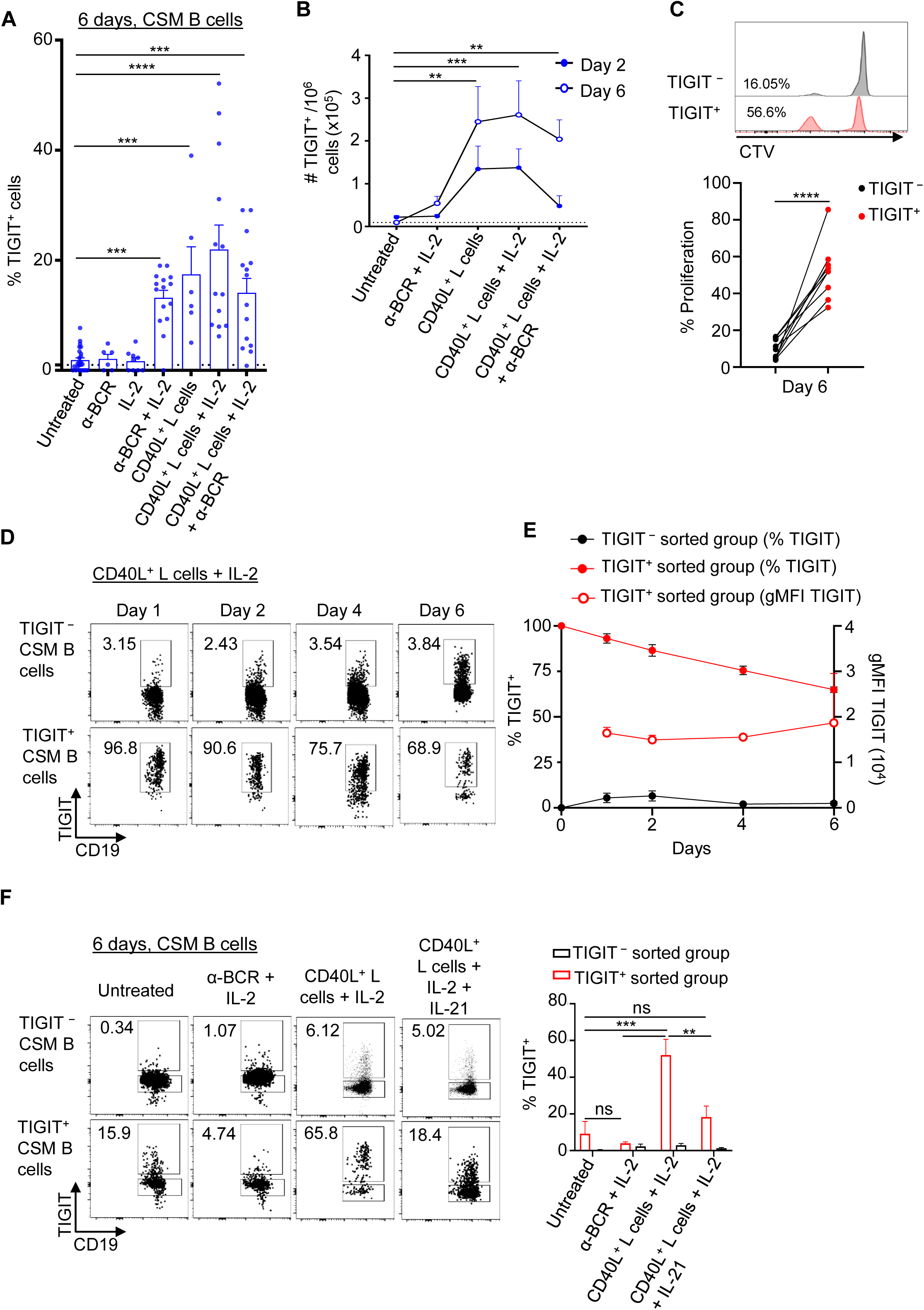
IL-2 and CD40L help maintain surface TIGIT expression on CSM B cells. **(A)** Purified CSM B cells were incubated for 6 days in the indicated conditions. The frequency of TIGIT^+^ CSM B cells was measured by flow cytometry. Individual dots represent data generated with cells from different healthy donors (N = 6-14). **(B)** Actual numbers of TIGIT^+^ CSM B cells in (**A**) were counted by flow cytometry using cell counting beads. **(C)** Purified CSM B cells were stained with CTV and cultured for 6 days in the presence of IL-2 and γ-irradiated CD40L-L cells. Proliferation of TIGIT^+^ and TIGIT^−^ B cell population was measured by flow cytometry. **(D)** Purified CSM B cells were incubated for 6 days in the presence of IL-2 and γ-irradiated CD40L-L cells and then FACS-sorted TIGIT^+^ and TIGIT^−^ CSM B cells were cultured for additional 6 days under similar condition. The frequency of TIGIT^+^ CSM B cells was measured at indicated days by flow cytometry. Representative FACS data are shown. **(E)** Summary of the frequency of TIGIT^+^ CSM B cells and geometric mean fluorescence intensity (gMFI) of surface TIGIT were measured by flow cytometry at indicated days. Cells from 4 healthy donors were tested. **(F)** FACS-sorted TIGIT^+^ and TIGIT^−^ CSM B cells as in **D** were cultured for additional 6 days under the indicated conditions. The frequency of TIGIT^+^ CSM B cells was measured by flow cytometry. Representative dot plots (**left**), and summarized data (**right**) are shown. Data are presented as mean ± SEM. Statistical analyses were performed using one-way ANOVA with Tukey’s multiple comparisons test (**A**), two-way ANOVA with Dunnett’s multiple comparisons test (**B**), or Tukey’s multiple comparisons test (**F**), and two-tailed unpaired t-test (**C**). ns, not significant, ∗ <0.05, ∗∗ <0.01, ∗∗∗ <0.001, ∗∗∗∗ <0.0001.

We next investigated the stability of surface TIGIT expressed on CSM B cells over time. After culturing CSM B cells for 6 days in the presence of IL-2 and CD40L, TIGIT^+^ and TIGIT^−^ CSM B cells were sorted by FACS. They were then cultured for additional 6 days in the presence of IL-2 and CD40L (**Figure 2D).** The frequency of CSM B cells expressing surface TIGIT gradually decreased, but the majority of CSM B cells, approximately 60-70%, maintained surface TIGIT expression at least by day 6 without decreasing surface TIGIT intensity (**Figure 2E**), indicating sustained expression of TIGIT at the single-cell level despite a reduction in the overall TIGIT^+^ B cell fraction. In contrast, TIGIT^−^ population failed to re-establish a TIGIT^+^ phenotype following re-stimulation (**Figures 2D and 2E**), suggesting that the acquisition and maintenance of TIGIT expression could be restricted to a subset of CSM B cells and might not be readily induced *de novo* under the conditions tested.

We next tested whether CD40L could contribute to the maintenance of surface TIGIT expressed on CSM B cells. In the absence of CD40L, TIGIT^+^ CSM B cells significantly lose surface TIGIT expression by day 6 (**Figure 2F**). Neither anti-BCR nor IL-2 could maintain surface TIGIT on CSM memory B cells. Of interest, addition of IL-21 significantly reduced surface TIGIT expression.

Collectively, IL-2 and CD40L play important roles in the generation of TIGIT^+^ CSM B cells and CD40L also contributes to the maintenance of surface TIGIT expressed on CSM B cells, supporting an important role for sustained T cell-derived signals in preserving this regulatory B-cell phenotype. Moreover, our data demonstrate that CD40L can promote CSM B cell viability after 6 days of culture (**Figure S6**), consistent with previous report,^46^ providing additional support for its role in sustaining TIGIT^+^ CSM B cells. Conversely, IL-21-induced reduction of surface TIGIT expression suggested that T cells, especially IL-21-producing Tfh cells, may counteract the IL-2/CD40L-mediated maintenance of TIGIT^+^ B cells *in vivo*.

### TIGIT⁺ CSM B cells acquire distinct immunoregulatory and migratory molecular signatures

To further define TIGIT^+^ CSM B cells generated with IL-2 and CD40L, a bulk RNA-seq analysis of FACS-sorted TIGIT^−^ and TIGIT^+^ CSM B cells was performed (**Figure 3**). Principal component analysis (PCA) revealed distinct clustering between TIGIT^+^ and TIGIT^−^ CSM B cell populations based on their transcriptomics profiles (**Figure 3A**). Differential gene expression analysis (p < 0.05, fold change > 2) identified a total of 470 differentially expressed genes (DEGs), with 325 downregulated and 145 upregulated genes in TIGIT⁺ CSM B cells compared to TIGIT⁻ CSM B cells (**Figures 3B and 3C**).

**Figure 3.**
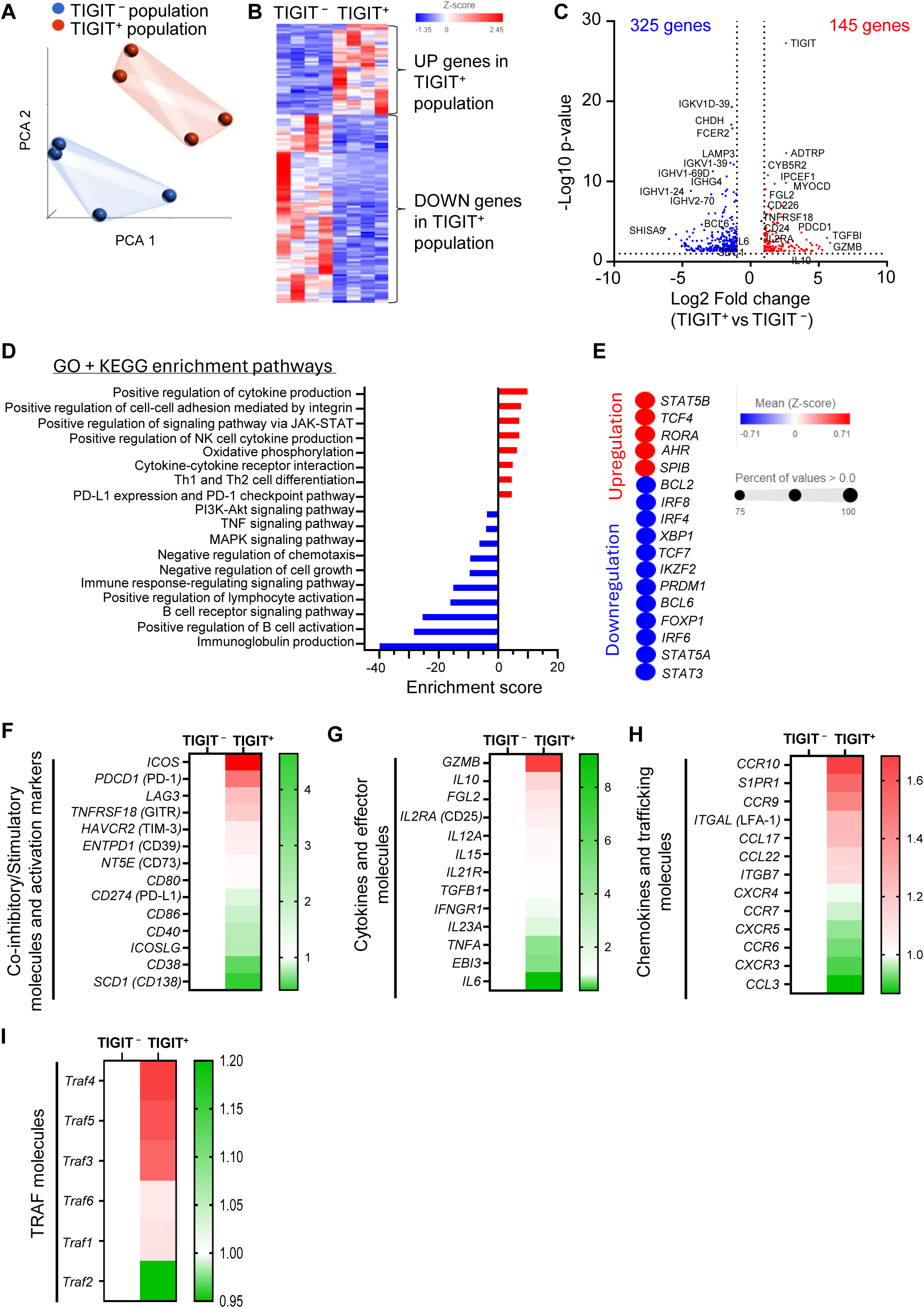
Molecular features of IL-2/CD40L-induced *ex vivo*-generated TIGIT⁺ CSM B cells. Purified CSM B cells from healthy donors (N = 4) were cultured for 6 days in the presence of IL-2 and γ-irradiated CD40L-L cells. TIGIT^+^ and TIGIT^−^ CSM B cells were FACS-sorted for RNA-seq analysis. **(A)** Principal component analysis (PCA) showing clustering of the 8 samples from 4 healthy donors. **(B)** Heatmap of 470 Differentially Expressed Genes (DEGs) (|log_2_ FC| > 0.5, p-value <0.05) in TIGIT^−^ and TIGIT^+^ populations. **(C)** Volcano plot (|log_2_ FC| > 0.5, p-value <0.05) depicting DEGs in TIGIT^+^ populations. Red dots represent up-regulated genes in TIGIT^+^ populations, and blue dots represent down-regulated genes in TIGIT^+^ populations. **(D)** KEGG and GO pathway enrichment analyses of DEGs. Red bars represent up-regulated pathways in TIGIT^+^ populations, and blue bars represent down-regulated pathways. **(E)** Dot plot showing transcription factors that are upregulated (red) or downregulated in TIGIT^+^ CSM B cells. (**F-I**) Heatmaps presenting expression levels of genes normalized on TIGIT^−^ CSM B cells for (**F**) co-inhibitory/stimulatory molecules and activation markers, (**G**) cytokines and other effector molecules, (**H**) chemokine/chemokine receptors and molecules related to tissue homing/retention, (**I**) TRAF molecules.

To gain insight into the biological pathways associated with TIGIT expression, we performed Kyoto Encyclopedia of Genes and Genomes (KEGG) and Gene Ontology (GO) pathway enrichment analyses. TIGIT^+^ CSM B cells were enriched for pathways involved in cytokine production, PD-L1/PD-1 checkpoint signaling, cell adhesion, and Th1/Th2 cell differentiation. In opposite, genes associated with lymphocyte activation, immune response, B cell activation, and Ig production were significantly downregulated, along with negative regulation of chemotaxis and cell growth (**Figure 3D**). Notably, JAK-STAT signaling pathway was enriched in TIGIT⁺ B cells, which is in line with prior data showing that STAT5 is critical for IL-2-induced TIGIT expression (**Figures 1F and 1G**). TIGIT^+^ CSM B cells also upregulate both *STAT5B* and *TCF-4* expression (**Figure 3E**). Additional transcription factors such as *RORA* and *AHR* that can induce IL-10 expression by B cells in mice,^47^ were also upregulated in TIGIT^+^ B cells, while B cells differentiation-related genes, *PRDM1* and *XBP1* were downregulated in TIGIT^+^ CSM B cells. *BCL6*, a master regulator of germinal center (GC) B cells, was also lower in TIGIT^+^ than TIGIT^−^ CSM B cells.

Further analyses of genes related to immune regulation revealed that TIGIT⁺ CSM B cells expressed increased levels of genes for PD-1, LAG-3, GITR, and ICOS along with moderate increase of TIM-3, CD39 and CD73. In contrast, they expressed decreased levels of genes associated with B cell activation markers, including CD86 and CD40, and differentiation markers such as CD38 and CD138 (**Figure 3F**), which was consistent with the downregulation of Ig heavy chain genes (**Figure 3C**). Additionally, TIGIT^+^ B cells also upregulated anti-inflammatory mediators such as *IL10* and *GZMB* genes, while pro-inflammatory cytokines *TNFA* and *IL6* genes were downregulated (**Figure 3G**).

Chemokine receptors expressed on lymphocytes contribute to their migration towards specific tissues or organs. Of interest, TIGIT^+^ B cells express increased levels of *CCR10*, *CCR9*, *ITGAL* and *ITGB7* (**Figure 3H**). CCR9 and β7 integrin contribute to lymphocyte homing to intestinal mucosa, while CCR10 is known to be a pan-mucosal homing receptor, although it also contributes to lymphocyte migration to skin.^48,49^ High expression of S1PR1, a migratory receptor,^50,51^ further supports the notion that TIGIT^+^ B cells have the capacity to migrate to peripheral tissues. TIGIT^+^ B cells also showed enrichment in *CCL17* and *CCL22* genes, chemokines involved in the recruitment of CCR4^+^ Th2-type T cells, including Th2 Tregs,^52–54^ suggesting a potential role in shaping local immune responses through cellular recruitment and polarization. Of note, genes of nearly all TRAF family members were also found to be more enriched in TIGIT^+^ B cells (**Figure 3I**), supporting sustained engagement of CD40-mediated signaling pathways and their downstream PI3K-AKT network.

Collectively, TIGIT^+^ CSM B cells generated with IL-2 and CD40L display unique molecular phenotypes, including increased expression of other inhibitory molecules and peripheral tissue homing receptors, that distinguish them from TIGIT^−^ CSM B cells. TIGIT^+^ CSM B cells also show significant downregulation of genes related to B cell differentiation and Ig secretion.

### Immunoregulatory and migratory properties of TIGIT⁺ CSM B cells

We further define TIGIT^+^ CSM B cells by measuring surface expression level of 21 different molecules that could be associated with their immune regulatory functions, and compared those expressed on TIGIT^−^ CSM B cells. **Figure 4A** shows a t-SNE plot clearly separating TIGIT^+^ from TIGIT^−^ CSM B cells based on the expression of surface proteins (**Figures 4B and S7**). Consistent with the transcriptomics data (**Figure 3F**), TIGIT^+^ CSM B cells express higher levels of PD-1, LAG3, GITR, and ICOS than TIGIT⁻ CSM B cells (**Figure 4B**). Interestingly, CTLA-4 and CD39 were also upregulated despite being undetectable and slightly increased in TIGIT^+^ CSM B cells, respectively, at the transcript level. In line with RNA-seq results, CD38 expression was significantly reduced, whereas the expression of CD40 was upregulated on TIGIT^+^ CSM B cells (**Figure 4B**). Cell surface expressions of CD73 and CD25 (IL2RA) also showed a trend toward upregulation on TIGIT^+^ CSM B cells, but the differences were not statistically significant (**Figure S7**).

**Figure 4.**
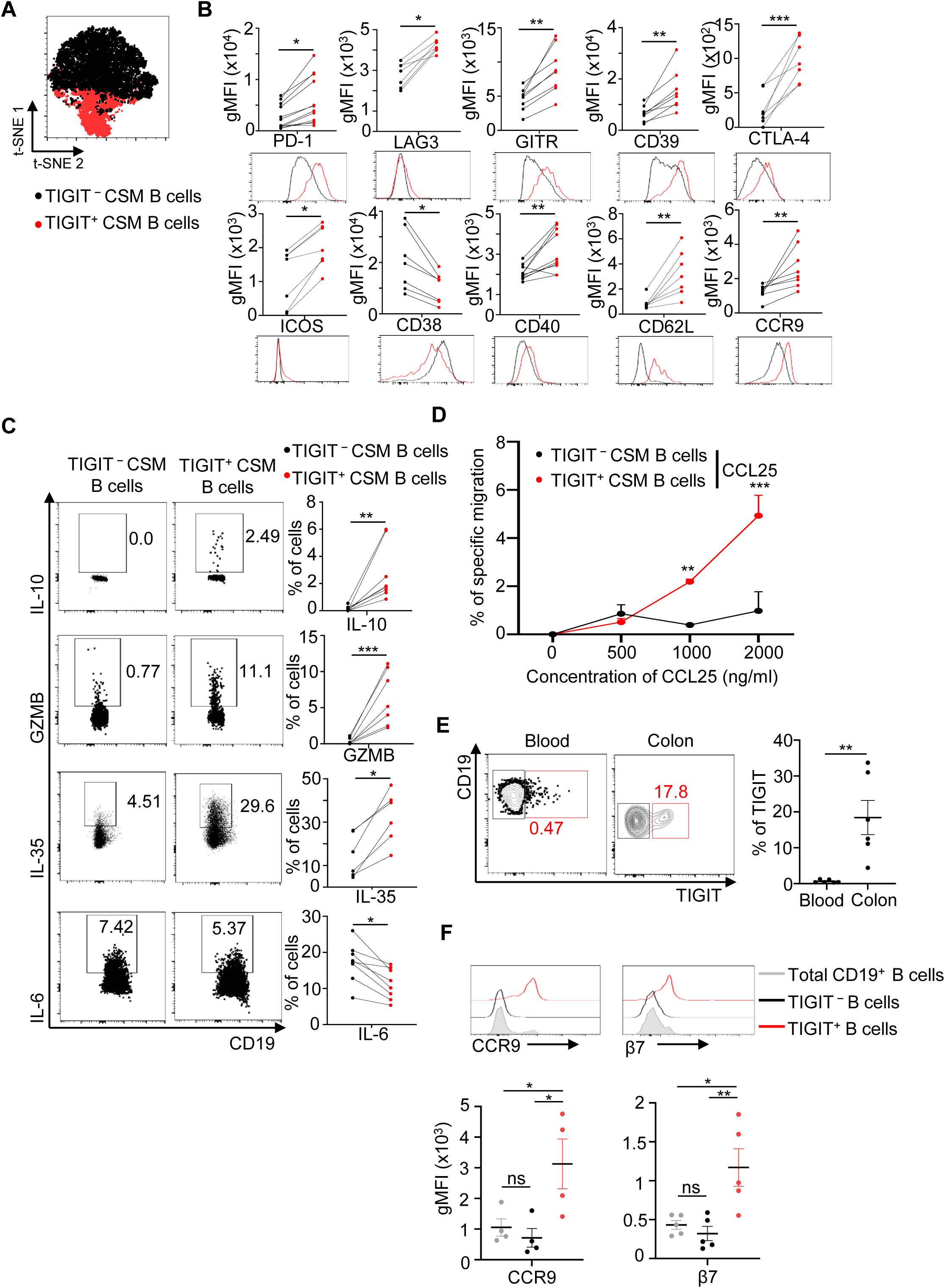
Immune regulatory phenotypes of IL-2/CD40L-induced *ex vivo*-generated TIGIT⁺ CSM B cells. Purified CSM B cells were cultured for 6 days in the presence of IL-2 and γ-irradiated CD40L-L cells. (**A**) A t-SNE plot showing the distribution of TIGIT^+^ and TIGIT^−^ CSM B cells from concatenated flow cytometry samples based on the expression profiles of 21 molecules (Figure 4B and **Figure S7**). (**B**) Surface expression levels of indicated molecules on TIGIT^+^ and TIGIT^−^ CSM B cells were measured by flow cytometry. Cells from healthy donors (N = 6-12) were tested. Each dot represents data generated with cells from different donors. Representative histograms for each marker are shown below each summarized graph. (**C**) Representative FACS data (**left**) and summarized data (**right**) showing the percentages of TIGIT^+^ and TIGIT^−^ CSM B cells expressing intracellular IL-10, GZMB, IL-35, and IL-6 in response to PMA/ionomycin. Each dot represents data generated with cells from different donors. (**D**) Chemotaxis assay with TIGIT^+^ and TIGIT^−^ CSM B cells (N = 4). FACS-sorted TIGIT^+^ and TIGIT^−^ CSM B cells were added to the upper wells of transwell and indicated concentrations of CCL25 were added in the lower wells. After 2 hours, the number of cells that migrated into the lower wells was determined by flow cytometry using cell counting beads. (**E**) The frequency of TIGIT^+^ B cells in the blood and colon tissues was assessed by flow cytometry (**left**). Summarized data of 5 donors are shown (**right**). (**F**) Surface CCR9 and β7 expression levels on TIGIT^+^ and TIGIT^−^ B cells in colon tissues were measured by flow cytometry (**upper**). Summarized data of tissues from 5 donors (**lower**). Data are presented as mean ± SEM. Statistical significance was determined using a two-tailed unpaired t-test (**B, C, E**), two-way ANOVA with Tukey’s multiple-comparisons test (**D**), and one-way ANOVA with Tukey’s multiple-comparisons test (**F**). ns, not significant, ∗ <0.05, ∗∗ <0.01, ∗∗∗ <0.001,

Consistent with the data in **Figure 3G**, TIGIT^+^ CSM B cells also expressed increased IL-10, granzyme B, IL-35, while expressing decreased IL-6 in response to PMA/ionomycin (**Figure 4C**). These findings support the acquisition of a broad immunoregulatory program consistent with the transcriptomic profile identified by RNA-seq analysis.

Flow cytometric analysis also confirmed increased surface expression of CCR9 and also CD62L protein, critical for lymphocyte migration, on TIGIT^+^ CSM B cells (**Figure 4B**). CCR9 expressed on TIGIT^+^ CSM B cells were functional, as they were able to migrate toward CCL25 in a concentration-dependent manner, the cognate ligand for CCR9, in transwell migration assays (**Figure 4D**). Although both CCR10 and β7 integrin are also expressed, differences between TIGIT^+^ and TIGIT^−^ CSM B cells were not significant across donors (**Figure S7**). Expression levels of CCR6, CCR7, CXCR3, and CXCR5 were not significantly different between the two CSM B cell populations. The *in vivo* relevance of the increased expression of CCR9 on TIGIT^+^ CSM B cells was further confirmed with increased frequency of TIGIT^+^ B cells in human colon tissues (**Figure 4E**). TIGIT^+^ B cells in the colon tissues also express higher levels of both CCR9 and β7 integrin than TIGIT^−^ B cells (**Figure 4F**).

Therefore, we concluded that IL-2/CD40L-induced *ex vivo*-generated TIGIT^+^ CSM B cells possess not only surface TIGIT, but also additional immune regulatory features characterized by the upregulation of a variety of inhibitory receptors and soluble mediators. In addition, TIGIT^+^ CSM B cells also upregulate functional CCR9 expression, which is further supported by the presence of increased frequency of CCR9-expressing TIGIT^+^ B cells in human colon tissues, highlighting their physiological relevance *in vivo*.

### TIGIT^+^ CSM B cells show defect in total Ig secretion, but not IgA2

Based on the molecular (i.e. downregulation of Ig heavy chain genes and B cell differentiation markers) and cellular phenotypes (i.e., downregulation of surface CD38 expression), we further defined TIGIT^+^ CSM B cell surface Ig expression by measuring the amounts of total Ig as well as Ig isotypes secreted from TIGIT^+^ CSM B cells. These were then compared with those secreted by TIGIT^−^ CSM B cells.

Purified CSM B cells were cultured for 6 days in the presence of IL-2/CD40L and surface Ig staining was performed. As shown in **Figure 5A**, an decreased frequency of IgG^+^ B cells was observed in TIGIT^+^ CSM B cells, when compared to TIGIT^-^ B cells. However, the frequency of IgA^+^ B cells, especially IgA2^+^ B cells, was higher in TIGIT^+^ CSM B cells than TIGIT^-^ CSM B cells (**Figures 5A and 5B**).

**Figure 5.**
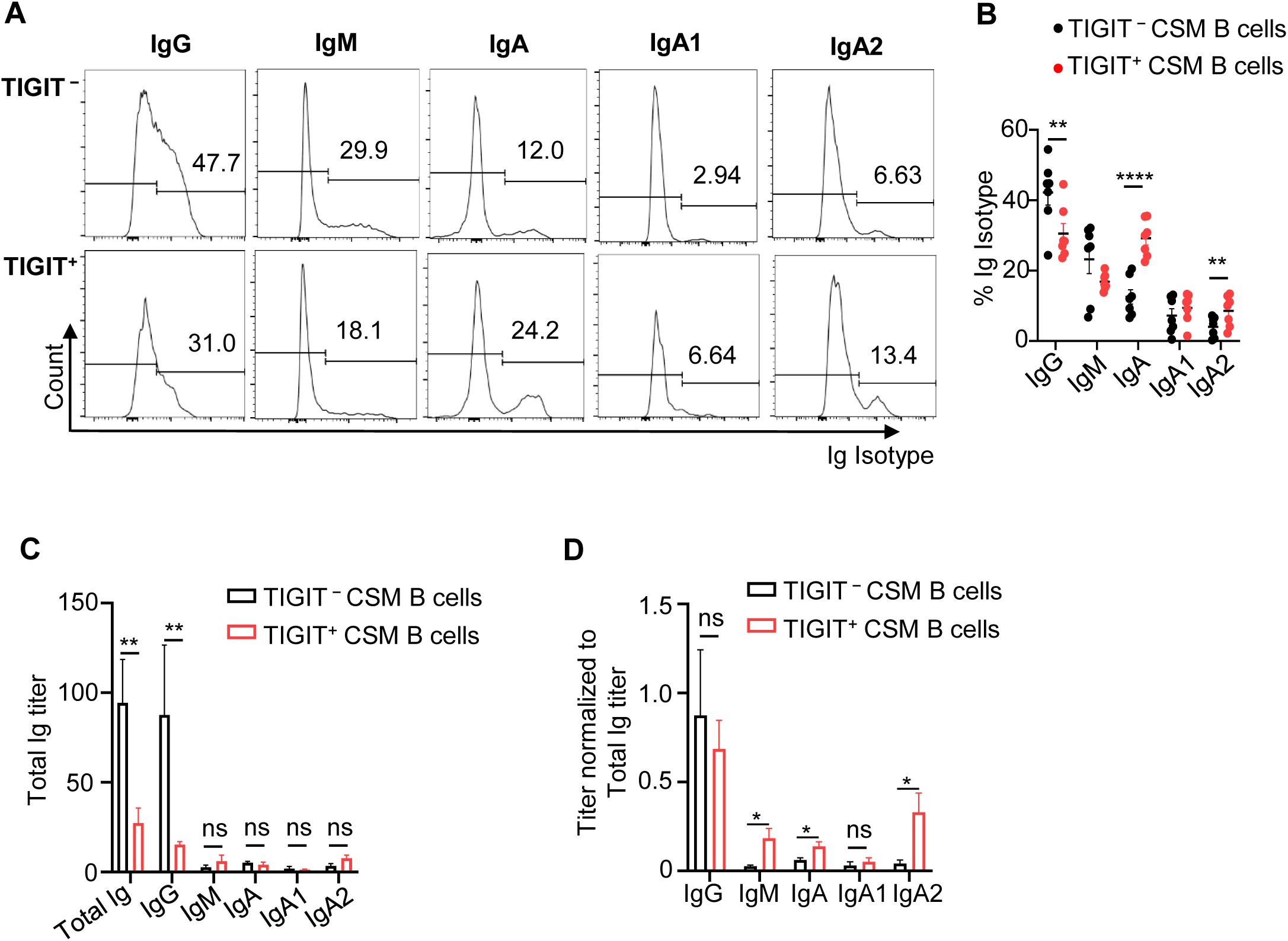
TIGIT^+^ CSM B cells secrete decreased amounts of total Igs, but not IgA2. (**A**) Purified CSM B cells were incubated for 6 days in the presence of IL-2 and γ-irradiated CD40L-L cells. Surface Igs were stained on TIGIT^−^ and TIGIT^+^ CSM B cells and analyzed by flow cytometry. Representative FACS data are presented. (**B**) Summarized data of (**A**) are presented. Individual dots represent data generated with cells from different healthy individuals (N = 7). (**C**) FACS-sorted TIGIT^+^ and TIGIT^−^ CSM B cells were cultured for additional 3 days in the presence of IL-2 and γ-irradiated CD40L-L cells. The amounts of Igs in the culture supernatants were measured by ELISA. (**D**) Data in (**C**) are expressed as titer of each isotype normalized to the titer of total Igs. Data in (**B**, **C**-**D**) are presented as mean ± SEM. Statistical significance was determined using a two-way ANOVA with Sidak’s multiple-comparisons test (**B, C-D**). ns, not significant, ∗ <0.05, ∗∗ <0.01, ∗∗∗∗ <0.0001.

We next measured the amount of total Ig and Ig isotypes secreted from FACS-sorted TIGIT^+^ and TIGIT^−^ CSM B cells. We found that TIGIT^+^ CSM B cells secreted markedly decreased amounts of total Ig as well as IgG isotype (**Figure 5C**). Of great interest, TIGIT^+^ CSM B cells, compared to TIGIT^−^ CSM B cells, secreted significantly increased ratios of both IgM and IgA especially IgA2 (**Figure 5D**).

Collectively, these data indicate that IL-2/CD40L-induced *ex vivo*-generated TIGIT^+^ CSM B cells have significant defects in total Ig and IgG secretion and this is in line with their molecular features, including decreased expressions of Ig heavy chain genes and genes associated with B cell differentiation, and cellular features, including decreased expression of CD38. However, TIGIT^+^ CSM B cells secrete greater ratio of IgA especially IgA2 which is further supported by their functional migration (**Figure 4D**) and enrichment in the human colon tissue with increased CCR9 expression (**Figures 4E and 4F**).

### TIGIT^+^ CSM B cells control DC activation and T cell responses

To validate the immune regulatory function of IL-2/CD40L-induced *ex vivo*-generated TIGIT^+^ CSM B cells, we first tested their ability to control DC activation. To this end, monocyte-derived DCs (MDDCs) were stimulated with *Escherichia. Coli (E. coli)* lipopolysaccharide (LPS) and cultured for two days in the presence of *ex vivo*-generated TIGIT^+^ or TIGIT⁻ CSM B cells. *E. coli* LPS-stimulated MDDCs alone served as control. MDDCs cocultured with TIGIT^+^ CSM B cells had reduced expression of maturation-associated surface molecules (CD40, CD80, ICOS-L, and CCR7), when compared with MDDCs cocultured with TIGIT^−^ B cells or MDDCs alone (**Figure 6A and Figure S8A**). TIGIT^+^ CSM B cells also reduced CD83 and CD86 expression levels in average mean fluorescence intensity (MFI) values. Anti-TIGIT neutralizing antibody partially restored DC maturation as indicated by a reduced percentage of suppression (**Figure 6B**), indicating that surface TIGIT expressed on CSM B cells contributes to the suppression of MDDC surface maturation.

**Figure 6.**
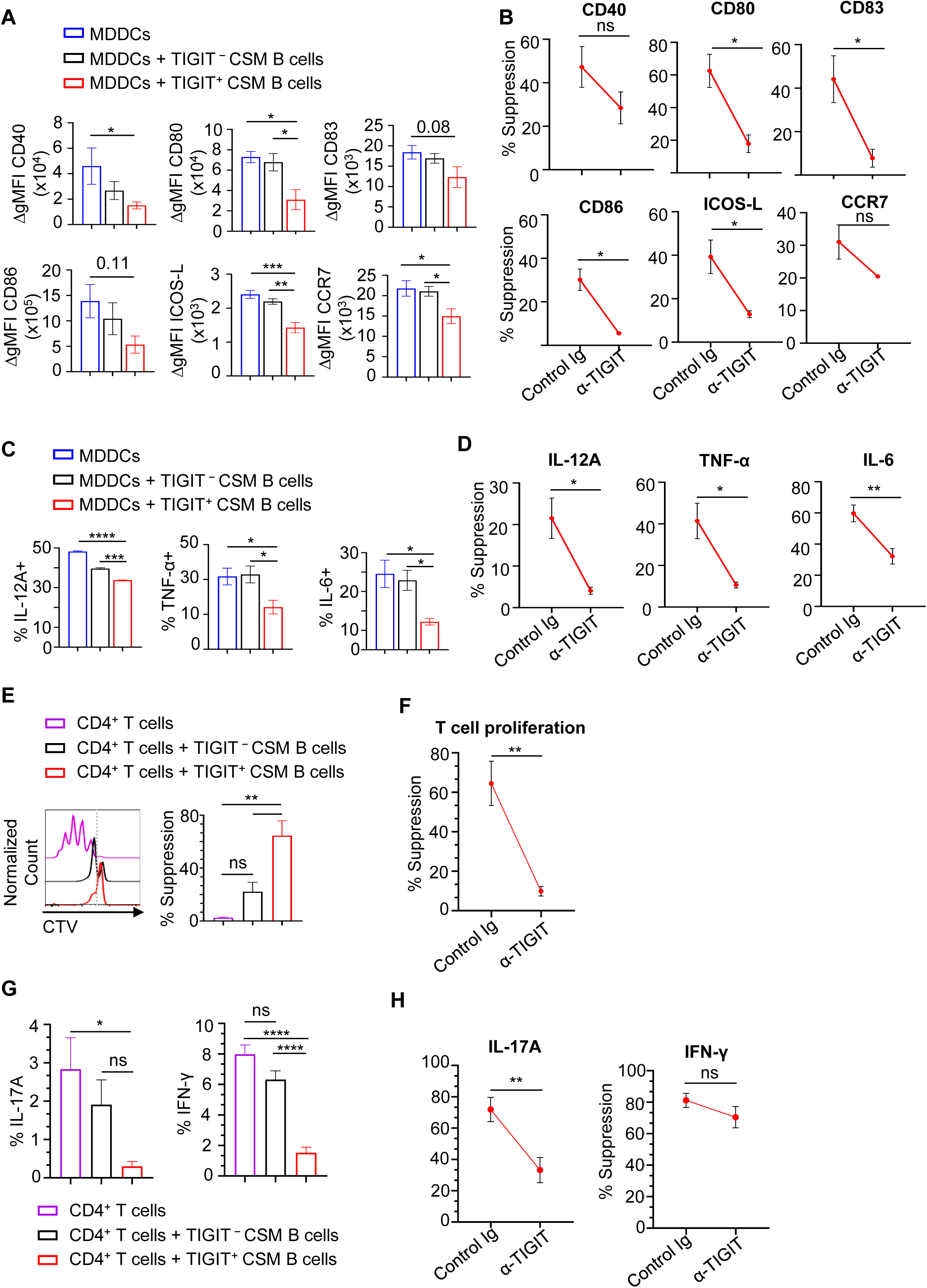
TIGIT⁺ CSM B cells can control MDDC activation and T cell responses. (**A-B**) *E. coli* LPS-stimulated MDDCs were co-cultured for 48 h with FACS-sorted TIGIT^+^ and TIGIT^−^ CSM B cells. Surface expression levels of CD40, CD80, CD83, CD86, ICOS-L, and CCR7 were measured by flow cytometry. Delta gMFI (ΔgMFI) values of individual molecules are presented (**A**). The same experiments were performed in the presence of control or anti-TIGIT neutralizing antibodies (**B**). Data are presented as the % suppression of individual molecules in each condition. (**C-D**) In (**A**), MDDCs were further stimulated for 5 h with PMA/ionomycin in the presence of brefeldin A before staining them with anti-IL-12A, anti-TNF-α, anti-IL-6 (**C**). The effects of anti-TIGIT neutralizing antibody on intracellular IL-12A, TNF-α, and IL-6 expression were assessed by calculating % suppression of individual cytokine expressions (**D**). (**E-F**) CTV-labeled naïve CD4^+^ T cells were stimulated with anti-CD3/28 dynabeads. After 6 days of co-culture with FACS-sorted TIGIT^+^ and TIGIT^−^ CSM B cells, CD4^+^ T cell proliferation was measured by flow cytometry and representative data (**left**) are presented. Summarized data (**right**, N = 4-8) are presented (**E**). The same experiments were performed in the presence of control and anti-TIGIT neutralizing antibodies. % suppression observed with anti-TIGIT antibody is presented (**F**). (**G-H**) From (**E**), CD4^+^ T cells were further stimulated for 5 h with PMA/ionomycin in the presence of brefeldin A before staining them with anti-IL-17A and anti-IFN-γ (**G**). The effects of anti-TIGIT neutralizing antibody on intracellular IL-17A and IFN-γ expression were assessed and % suppression of individual cytokine expressions are presented (**H**). Data are presented as mean ± SEM. Statistical analyses were performed using one-way ANOVA with Tukey’s multiple comparisons (**A, C, E, G**) or two-tailed unpaired t test (**B, D, F and H**). ns, not significant, ∗ <0.05, ∗∗ <0.01, ∗∗∗ <0.001, and ∗∗∗∗ <0.0001.

In addition, TIGIT^+^ B cells also suppressed IL-12A, TNF-α and IL-6 by LPS-stimulated MDDCs (**Figure 6C and Figure S8B**). Anti-TIGIT neutralizing antibody also partly recovered proinflammatory cytokine expression by the same LPS-activated MDDCs (**Figure 6D**).

We next investigated whether TIGIT^+^ CSM B cells could also directly control T cell responses. CellTrace Violet (CTV)-labeled naïve CD4^+^ T cells were stimulated with anti-CD3/28 dynabeads and then co-cultured with either TIGIT^+^ or TIGIT^−^ CSM B cells for six days. TIGIT^+^ B cells suppressed T cell proliferation substantially more effectively than TIGIT^−^ B cells, achieving approximately 65% inhibition at a 1:1 B : T cell ratio compared with approximately 20% inhibition by TIGIT^−^ B cells (**Figure 6E**). Blocking TIGIT with anti-TIGIT antibody resulted in the recovery of CD4^+^ T cell proliferation (**Figure 6F**). TIGIT^+^ CSM B cells also decreased IFN-γ and IL-17A expression by CD4^+^ T cells (**Figure 6G and Figure S8C**). The reduction in IL-17A expression was partially restored by neutralizing TIGIT, whereas IFN-γ expression showed only a modest decrease that did not reach statistical significance (**Figure 6H**).

Together, these results demonstrate that IL-2/CD40L-induced *ex vivo*-generated TIGIT^+^ CSM B cells can control DC activation and T cell responses, and TIGIT expressed on the CSM B cells plays an important role in controlling DC maturation and T cell responses.

### TIGIT^+^ CSM B cells ameliorate aGVHD with the enhancement of IL-10-producing Treg response

To investigate how TIGIT^+^ CSM B cells exert immunoregulatory activity *in vivo*, we employed a xenogeneic aGVHD model using NOG mice, in which human T cells are the major causes of the disease.

A single intravenous (i.v.) dose of FACS-sorted 4 × 10^5^ CSM B cells (containing ∼3.9 × 10^4^ TIGIT^+^ B cells, measured by FACS) at the time of PBMC (10 × 10^6^) transplantation inhibited aGVHD progression overtime (**Figure 7A**). Such inhibition was not observed when animals received TIGIT^+^ B cell-depleted CSM B cells, suggesting that TIGIT expression on CSM B cells contribute to the suppression of aGVHD progression.

**Figure 7.**
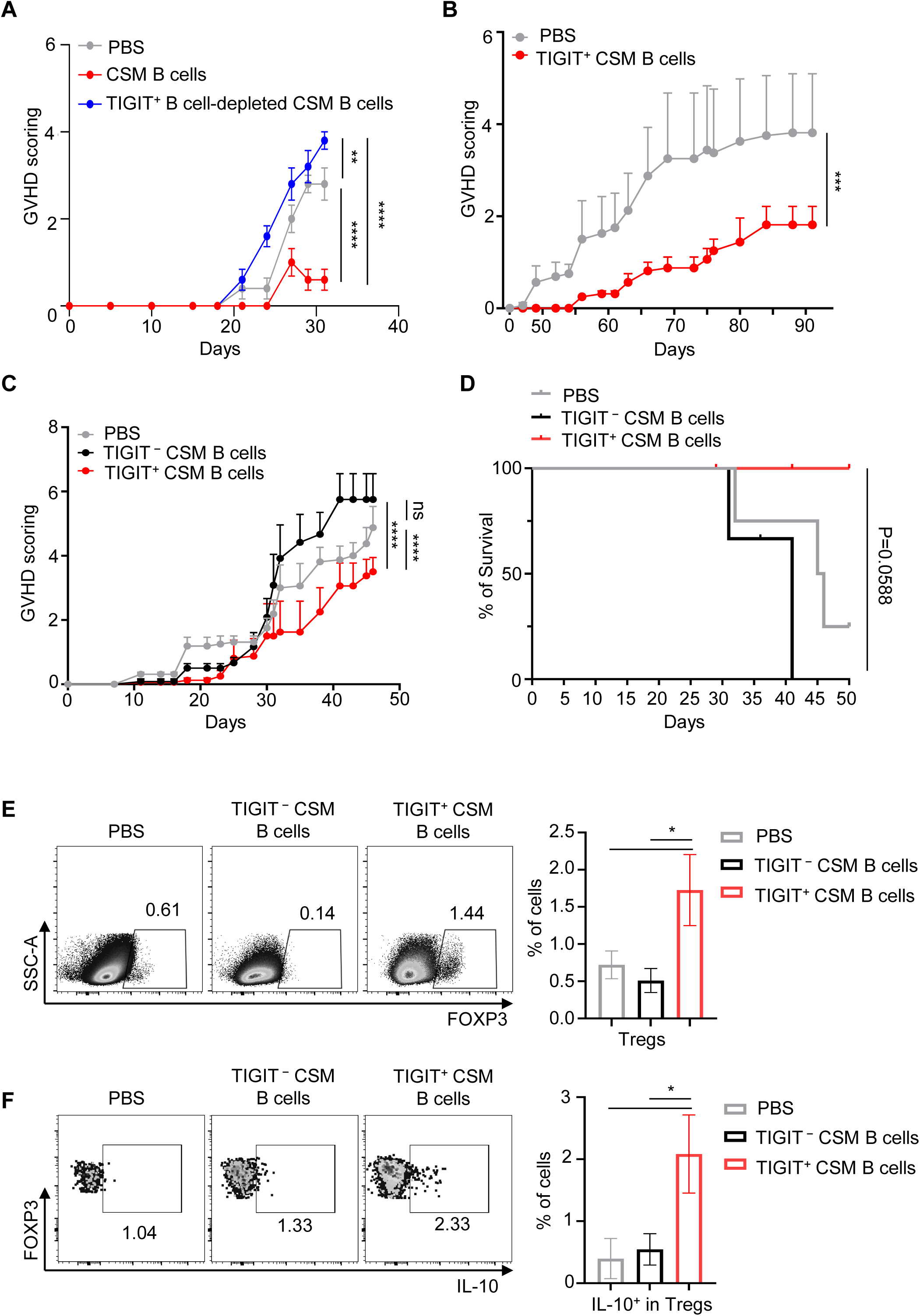
TIGIT⁺ CSM B cells suppress aGVHD progression with the enhancement of IL-10-producing Treg response. All NOG animals (4 animals per group) received PBMC (10 x 10^6^) via tail vein on day 0. (**A**) Animals were divided into 3 groups and received PBS, 4 x 10^5^ CSM B cells, or 4 x 10^5^ TIGIT^+^ B cell-depleted CSM B cells on day 0. GVHD disease scores were assessed. (**B**) Two groups of animals received PBS alone and FACS-sorted IL-2/CD40L-induced ex vivo-generated TIGIT^+^ CSM B cells (2 x 10^5^ cells), respectively, via tail vein on day 0. (**C-D**) Three groups of animals received PBS, TIGIT^−^ CSM B cells (7 x 10^4^ cells), and TIGIT^+^ CSM B cells (7 x 10^4^ cells), respectively, via tail vein on day 0. GVHD scores (**C**) and % survival (**D**) were assessed. (**E-F**) Splenocytes from experiment in (**C-D**) were stained for CD4^+^FOXP3^+^ Tregs (**E**) and IL-10^+^ in Tregs (**F**). Data are presented as mean ± SEM. Statistical analyses were performed using two-way ANOVA with Tukey’s (**A-C**) or one-way ANOVA with Tukey’s multiple comparisons (**E-F**). Statistical significance for Kaplan–Meier survival curve (**D**) was determined using the log-rank (Mantel– Cox) test. ∗ <0.05, ∗∗ <0.01, ∗∗∗ <0.001, ∗∗∗∗ <0.0001.

We next assessed the immunosuppressive activity of IL-2/CD40L-induced *ex vivo*-generated TIGIT^+^ CSM B cells. Adoptive transfer of 2 × 10^5^ TIGIT^+^ CSM B cells together with 10 × 10^6^ human PBMCs significantly attenuated disease progression compared with PBS-treated control (**Figure 7B**). To further validate the immune regulatory function of TIGIT^+^ CSM B cells, we next performed the same experiments with reduced numbers of *ex vivo*-generated TIGIT^+^ CSM B cells (7 × 10^4^). Remarkably, adoptive transfer of only 7 × 10^4^ TIGIT^+^ CSM B cells also significantly delayed and reduced aGVHD progression when compared with the two control treatments, PBS and TIGIT^−^ CSM B cells (**Figure 7C**). Such delayed progression of aGVHD was accompanied by improved animal survival (**Figure 7D**), highlighting the role of TIGIT^+^ CSM B cells in amelioration of aGVHD. aGVHD scores of individual animals in the three groups are presented in **Figure S9A**, showing that three of four mice receiving TIGIT^+^ CSM B cells maintained GVHD scores below 4 at 50 days, whereas only one animal in the PBS group and none in the TIGIT^−^ CSM B cell group achieved comparable disease control.

To investigate the mechanisms underlying the TIGIT^+^ CSM B cell-mediated suppression of aGVHD progression, spleens and tissues, including livers and intestines, were collected at sacrifice for flow cytometric and histological analyses, respectively. Although there was a trend toward decreased frequency of IL-17A^+^ CD4^+^ T cells in animals treated with TIGIT^+^ CSM B cells, no significant differences were observed in the frequencies of IFN-γ-, IL-17A-, or TNF-α-producing CD4^+^ or CD8^+^ T cells within the spleen (**Figure S9B**). Likewise, the frequency of IL-10^+^ CD4^+^ T cells was not significantly altered, although CD8^+^ T cells showed a trend toward increased IL-10 expression in animals who received TIGIT^+^ CSM B cells. However, we found that animals treated with TIGIT^+^ CSM B cells had significantly increased frequencies of CD4^+^FOXP3^+^ Tregs when compared to the other two groups of animals (**Figure 7E**). Furthermore, Tregs in the animals treated with TIGIT^+^ CSM B cells readily express IL-10 (**Figure 7F**), suggesting their immune regulatory function *in vivo*. Although CTLA-4 expression on Tregs in animals treated with TIGIT^+^ CSM B cells was not significantly increased, a consistent upward trend was observed (**Figure S9C**).

Therefore, we concluded that *ex vivo*-generated TIGIT^+^ CSM B cells can suppress human T cell-mediated inflammatory diseases, aGVHD in this case, with the increase of IL-10^+^ Tregs, supporting a novel TIGIT^+^ CSM B cell-initiated immune regulatory network that can be reinforced by the enhancement of Treg responses *in vivo*.

## DISCUSSION

This study investigated how B cells become immunosuppressive by identifying key factors governing molecular and cellular features of immunosuppressive B cells. Of great interest, CD4^+^ helper T cell-derived factors, IL-2 and CD40L, play critical roles in reprogramming CSM B cells to be immunosuppressive by the induction of not only TIGIT, but also a variety of currently known B cell-derived inhibitory molecules. With the acquisition of such broad immune regulatory phenotypes, TIGIT^+^ CSM B cells have significant defects in total Ig secretions, but they still secrete IgA2, suggesting their roles in the maintenance of mucosal immune homeostasis. TIGIT^+^ CSM B cell-mediated immune regulatory network is further reinforced by their ability to promote functional Treg cell response.

IL-2 was first discovered as a human T cell growth factor.^55^ It is produced primarily by CD4^+^ T cells,^56^ helps CD8^+^ T cell proliferation and activation, and has been used in the immunotherapy of certain types of cancers.^57^ In addition, IL-2 is also known to contribute to CD4^+^ T cell differentiation, especially Treg cell differentiation,^58^ expansion,^59^ and their regulatory functions.^60,61^ This study extends our knowledge on the immune regulatory function of IL-2 by demonstrating that it can also promote immune regulatory function of B cells by the induction of TIGIT expression on CSM B cells followed by the expression of a variety of inhibitory molecules. While IL-2-mediated STAT5 activation has been extensively studied in T cells as it contributes to regulatory and effector T cell differentiation,^43,62^ its role in governing immune regulatory function of B cells has not been previously studied. Concomitant upregulation of TIGIT and other inhibitory molecules, including IL-10, with the enhanced immune regulatory function of TIGIT^+^ CSM B cells might be associated at least in part with B cell-intrinsic IL-2 signaling that can promote IL-10 expression and regulatory functions as previously reported.^44^ The enrichment of STAT5B transcript in TIGIT⁺ B cells also supports sustained involvement of this particular signaling pathway beyond initial induction of TIGIT expression. In parallel, inhibition of PI3K or AKT also reduced TIGIT expression, implicating the roles of CD40L in the establishment of the regulatory phenotypes in CSM B cells. CD40 stimulation is known to induce IL-10 expression by B cells.^1^ Furthermore, a trend toward increased expression of multiple TRAF family members in TIGIT^+^ CSM B cells also supports molecular link between CD40 engagement and the PI3K-AKT activation pathway^63^ for not only TIGIT induction, but also the maintenance of immune regulatory phenotype of TIGIT^+^ CSM B cells. Notably, TIGIT expression was not readily induced *de novo* in sorted TIGIT⁻ CSM B cells, whereas established TIGIT⁺ CSM B cells largely maintained surface TIGIT expression following re-stimulation despite a gradual decline over time, potentially reflecting progressive cellular exhaustion. Further studies will be required to determine the mechanisms underlying this decline. Together with their distinct transcriptomic and phenotypic profiles, these findings support the concept that TIGIT⁺ CSM B cells represent a specialized regulatory subset rather than a transient activation state that can be acquired by all memory B cells. Collectively, these results highlight that IL-2 and CD40L not only generate TIGIT⁺ CSM B cells but also sustain their immunoregulatory phenotype.

It was also important to note that IL-21, another common γ-chain cytokine, did not promote the induction of TIGIT expression, but it significantly downregulated surface TIGIT expression on CSM B cells. In molecular levels, IL-21-induced STAT3 activation could suppress β-catenin and TCF-4 transcriptional activity that can be triggered by CD40L,^64^ which might lead to the downregulation of surface TIGIT expression, but more investigations are needed. More importantly, however, this IL-21-mediated downregulation of surface TIGIT expression supports another novel role of Tfh cells in antibody responses. Although additional studies are warranted, Tfh cells can presumably promote antibody responses by downregulating TIGIT expression on CSM B cells, as we observed TIGIT^+^ CSM B cells have significant defects in Ig secretion.

In line with transcriptomic profiles, IL-2/CD40-induced *ex vivo*-generated TIGIT⁺ CSM B cells upregulate not only surface inhibitory receptors, including PD-1, GITR, LAG-3, CD39, and CTLA-4, but also intracellular expressions of other effector molecules, including IL-10, IL-35, and granzyme B. Many of these molecules are traditionally associated with suppressive function of Tregs,^65–67^ suggesting convergence of regulatory programs across lymphocyte lineages. The discordance between CTLA-4 transcript and protein expression is consistent with extensive post-transcriptional regulation described in Tregs.^67^ Functionally, TIGIT⁺ CSM B cells efficiently suppressed DC maturation and proinflammatory cytokine expression as well as CD4⁺ T cell proliferation and inflammatory cytokine expression. Although TIGIT blockade partially reversed these effects, suppression caused by TIGIT^+^ CSM B cells was not completely abolished, indicating that other inhibitory molecules expressed by TIGIT^+^ CSM B cells contribute to their immune regulatory function, confirming that TIGIT^+^ CSM B cells are equipped with a multilayered-suppressive mechanism.^1,12,21,68,69^ Importantly, TIGIT^+^ CSM B cells were also capable of building a potent and prolonged immune regulatory network by the enhancement of Treg response. This was demonstrated in a xenogeneic mouse model of aGVHD, as the effect of TIGIT^+^ CSM B cell treatment lasted for longer than 6-12 weeks when there was no human B cell engraftment in any tissues. Mechanistically, administration of TIGIT⁺ CSM B cells promoted the expansion of IL-10-producing Tregs, providing a potential explanation for the prolonged regulatory effects observed *in vivo*.

TIGIT^+^ CSM B cells also display a unique mucosal regulatory phenotype. They express β7 integrin with significantly increased expression of functional CCR9, as demonstrated in migration assays in response to CCL25. They have significant defects in total Ig and IgG secretion, but not IgA2. In line with these observations, increased expression of CCR9 and IgA2 secretion, TIGIT⁺CCR9⁺ B cells were found to be enriched within human colon tissues. CCR9-CCL25 interactions are central to gut homing of lymphocytes.^48^ The preferential secretion of IgA2 is particularly notable given the specialized role of this isotype in intestinal immune homeostasis.^70–72^ In addition to CCR9, upregulation of *CCR10* and *S1PR1* mRNA expressions and CD62L protein further support a migratory program compatible with trafficking to peripheral mucosal tissues. Together, these findings suggest that TIGIT⁺ CSM B cells may be adapted for immune regulation at mucosal surfaces, where continuous exposure to dietary antigens and commensal microbiota requires tightly controlled immune responses. These phenotypes of TIGIT^+^ CSM B cells also share some features of mucosal B cells and tolerogenic IgA-producing plasma cells described in the gastrointestinal tract,^72–75^ raising the possibility that TIGIT⁺ CSM B cells contribute to maintenance of intestinal immune tolerance. Under steady-state conditions, the intestinal mucosa contains abundant CD4^+^ Th17 cells, and TIGIT expression in CSM B cells preferentially suppressed IL-17A over IFN-γ production by T cells, suggesting that TIGIT^+^ CSM B cells may be particularly effective at regulating Th17-dominated immune responses. Finally, the trend toward increased expression of *CCL17* and *CCL22* suggests that these cells also contribute to the recruitment of Th2-type Tregs at local tissues.

In summary, this study reinforces the emerging roles of B cells, especially TIGIT^+^ CSM B cells, in the suppression of inflammation and the maintenance of immune homeostasis.^2,21,69,76–78^ Such novel molecular and cellular features of TIGIT^+^ CSM B cells are programmed by the actions of IL-2 and CD40L that can be provided by CD4^+^ helper T cells. Once programmed, they can reinforce the immune regulatory network by the enhancement of Treg responses. Our study also supports the clinical development of TIGIT^+^ B cell-based therapy in the treatment of certain types of inflammatory diseases. Future studies might include the investigation of TIGIT intrinsic function within TIGIT^+^ CSM B cells, factors and physiologic conditions affecting surface TIGIT expression levels on CSM B cells and their associations with any disease phenotypes, and extended application of such novel immune regulatory function of TIGIT^+^ CSM B cells in any inflammatory disease settings.

### Limitations of the study

This study has several limitations. Although we identified IL-2 and CD40L as key factors that induce immunosuppressive TIGIT⁺ CSM B cells, the physiological conditions regulating their generation and maintenance in humans remain to be fully defined. Our findings are based primarily on *ex vivo* cultures of human B cells and a xenogeneic mouse model of aGVHD, which may not fully recapitulate the complexity of human inflammatory diseases. In addition, while TIGIT blockade partially reversed the suppressive activity of TIGIT⁺ CSM B cells, the relative contributions of other inhibitory molecules expressed by these cells were not fully dissected. Future studies using disease-specific models and genetic approaches will be important to define the intrinsic role of TIGIT in B cells and to evaluate the therapeutic potential of TIGIT⁺ CSM B cells across diverse inflammatory settings.

## RESOURCE AVAILABILITY

### Lead contact

Requests for further information and resources should be directed to the lead contact, SangKon Oh.

## Materials availability

This study did not generate new reagents.

## Data and code availability

- The sequencing data have been deposited to GEO. Accession number is listed in the key resources table.
- This paper does not report original code.

## ACKNOWLEDGEMENTS

The authors thank Dr. Justin Brady in Mayo Clinic Comprehensive Cancer Center for helping with the collection of the human colon tissues for the study. This project was supported (to S.O.) by the Mayo Clinic Transplantation Center, Mayo Clinic Ventures, and Mayo Clinic Comprehensive Cancer Center. This project was also supported by National Institutes of Health, National Institute of Allergy and Infectious Diseases (1R21AI190763-01A1 to S.O.)

## AUTHOR CONTRIBUTIONS

Conceptualization, Interpretation and Writing by L.B. and S.O.; Data analysis by L.B., A.C., and S.O.; Data visualization, L.B. and A.C.; Conducting experiments, L.B., A.C., A.Y., Resources, S.C., and S.B. All authors reviewed and approved the manuscript.

## STAR METHODS

### KEY RESOURCES TABLE

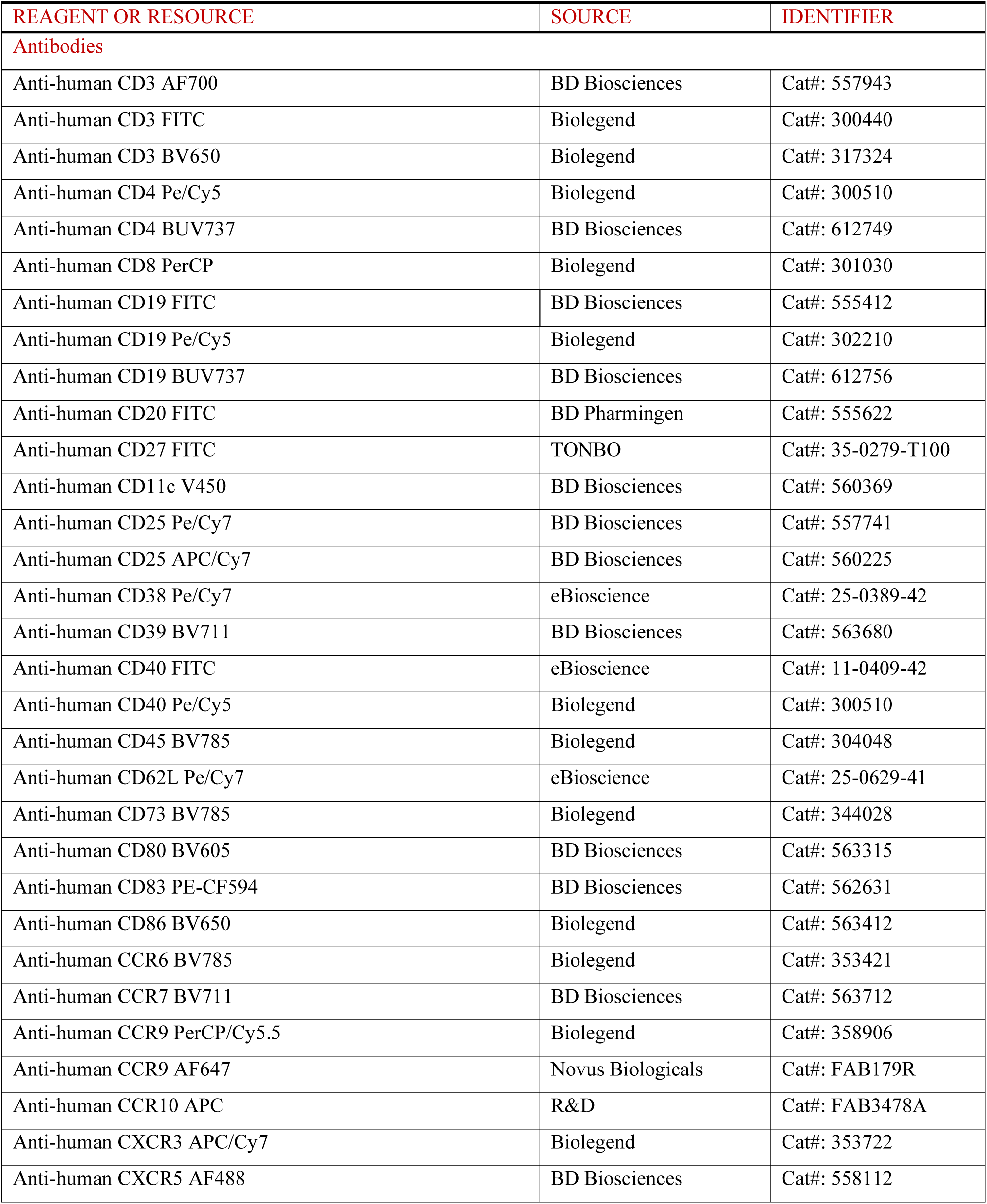
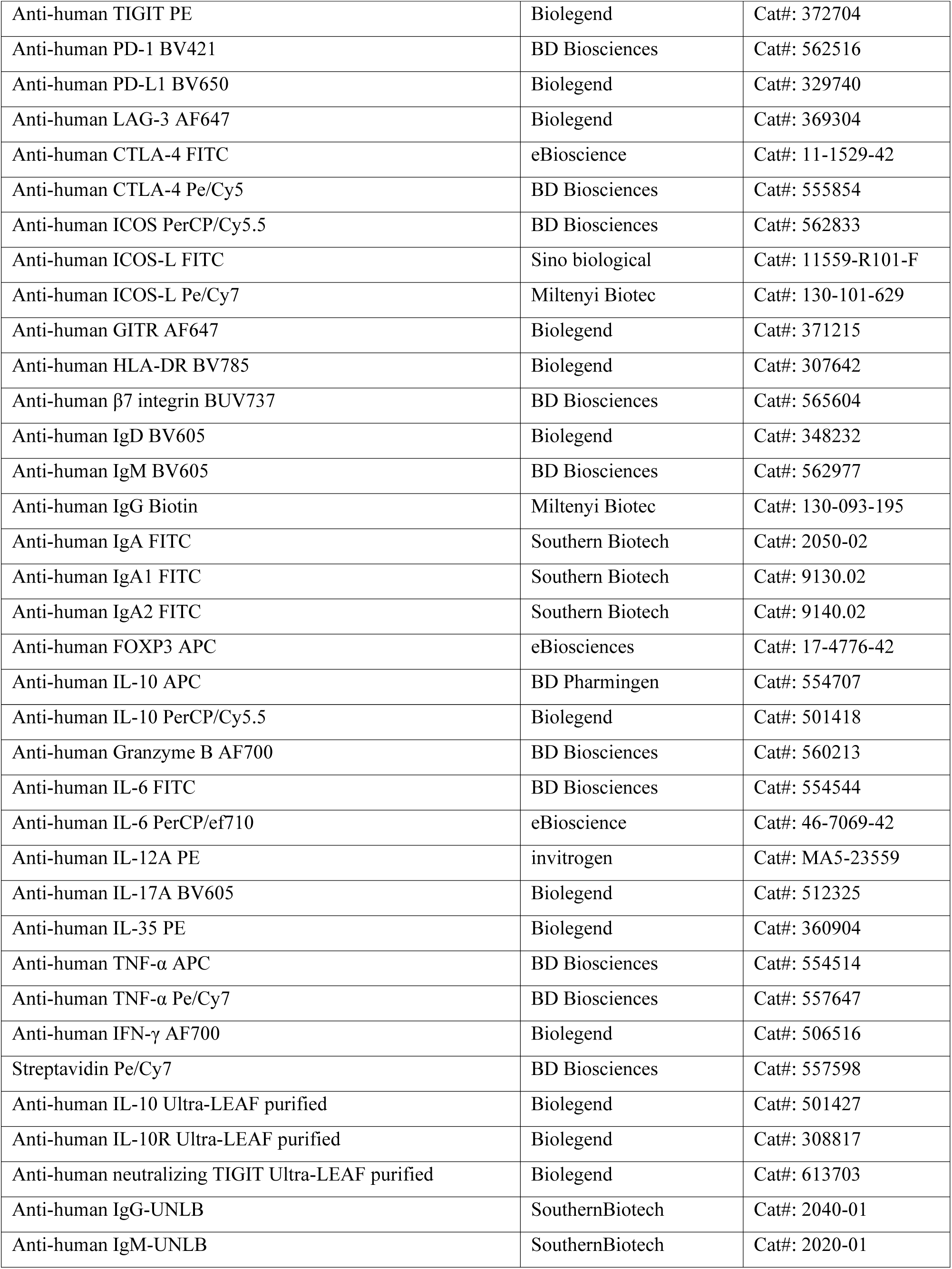
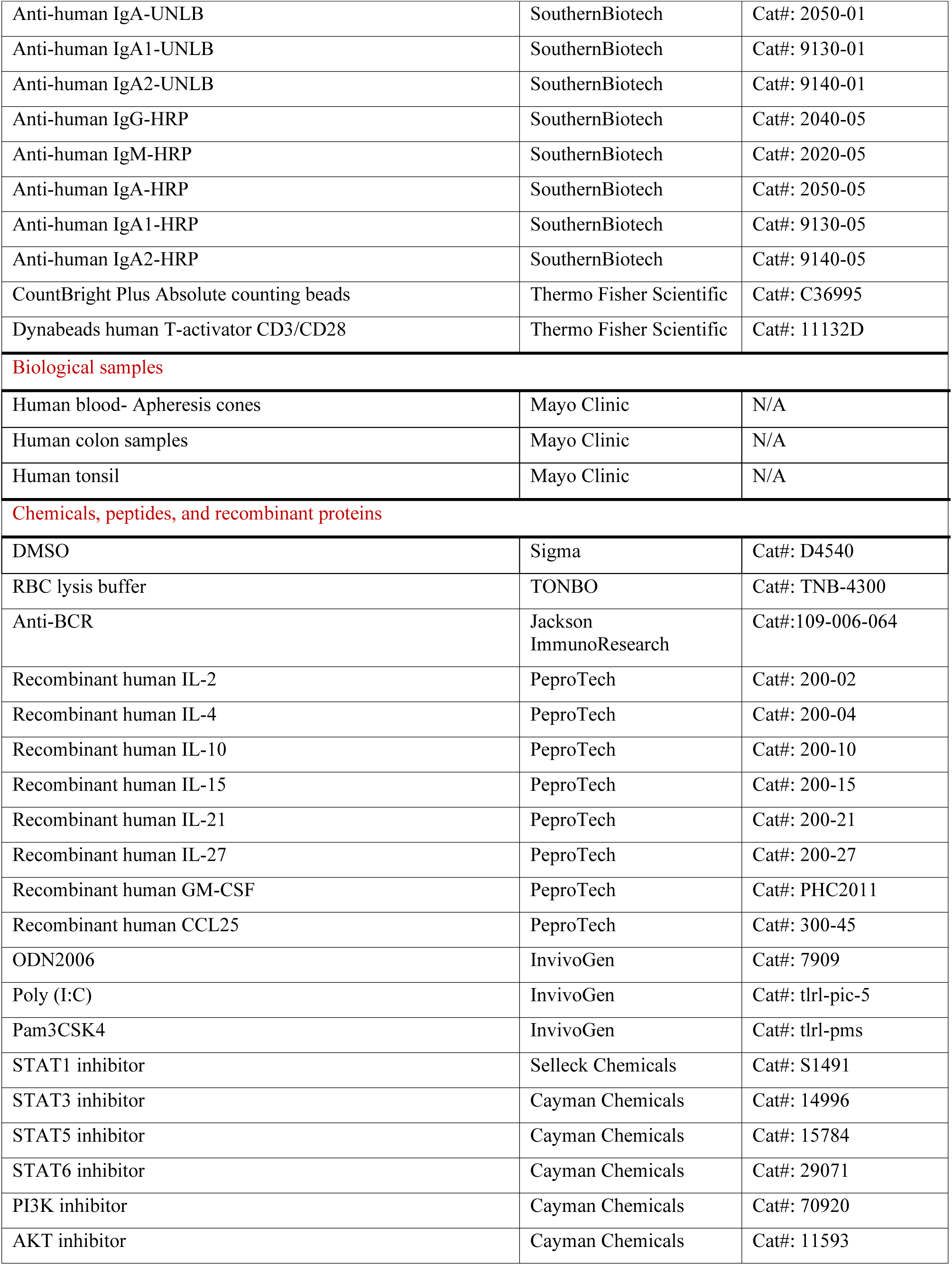
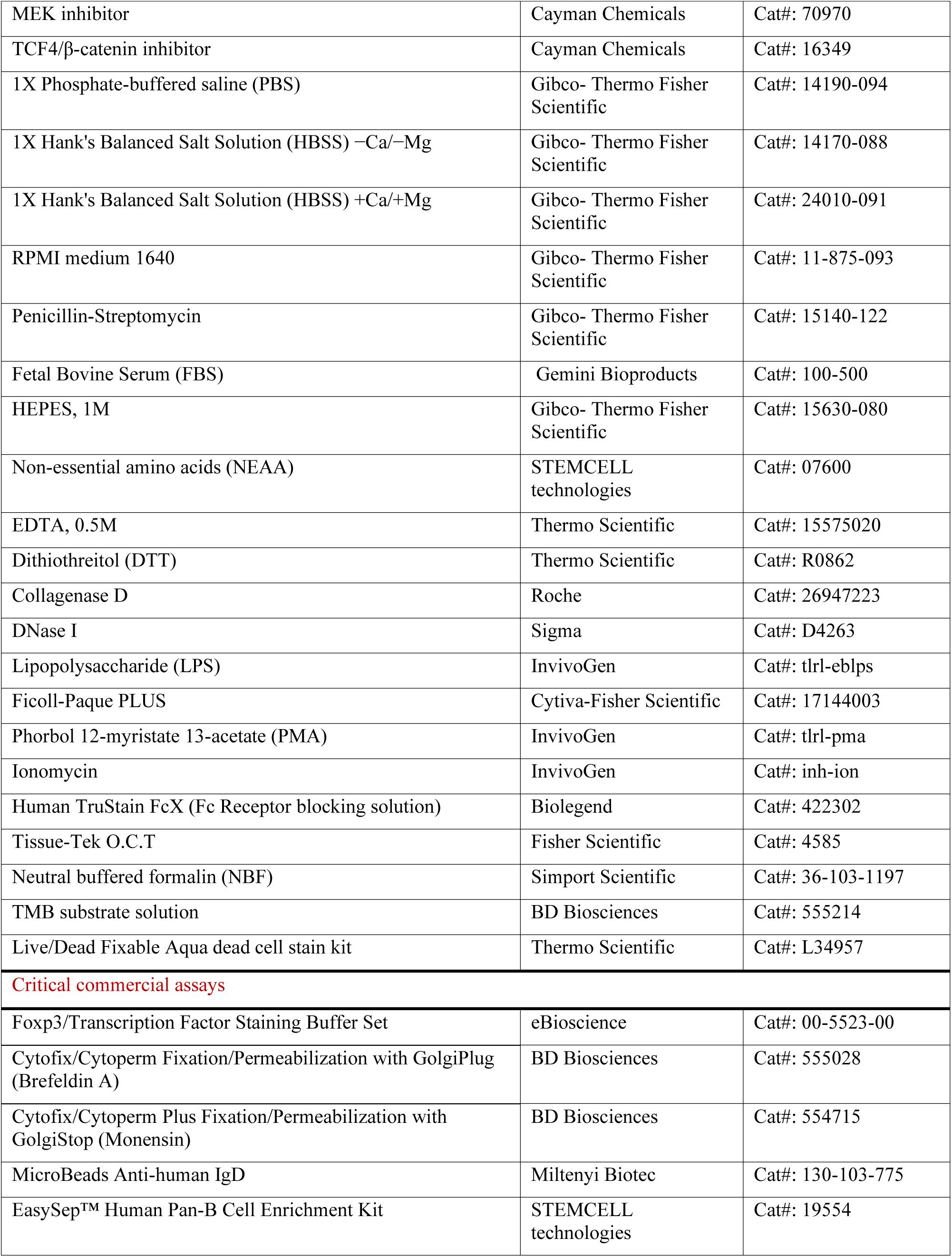
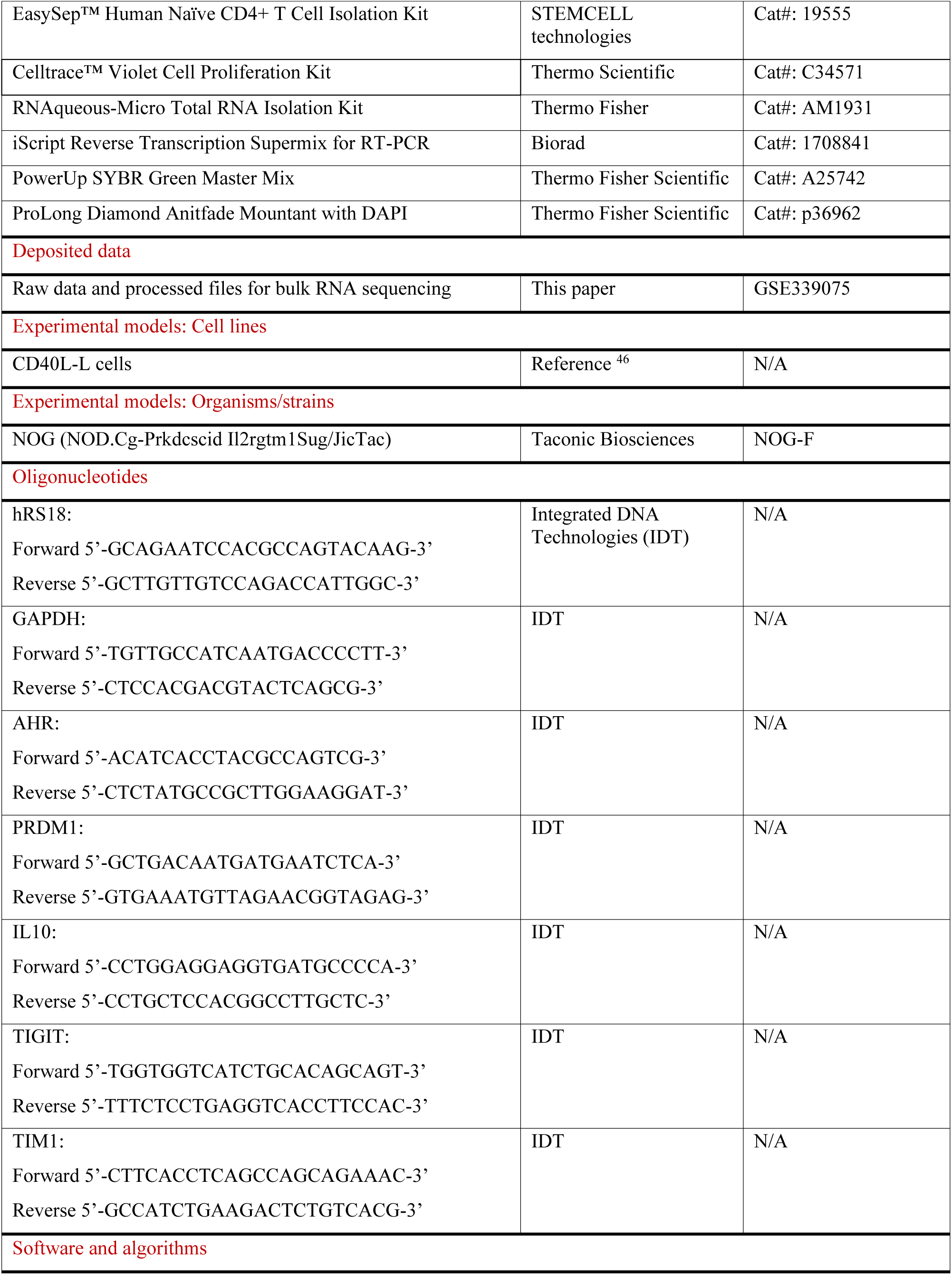
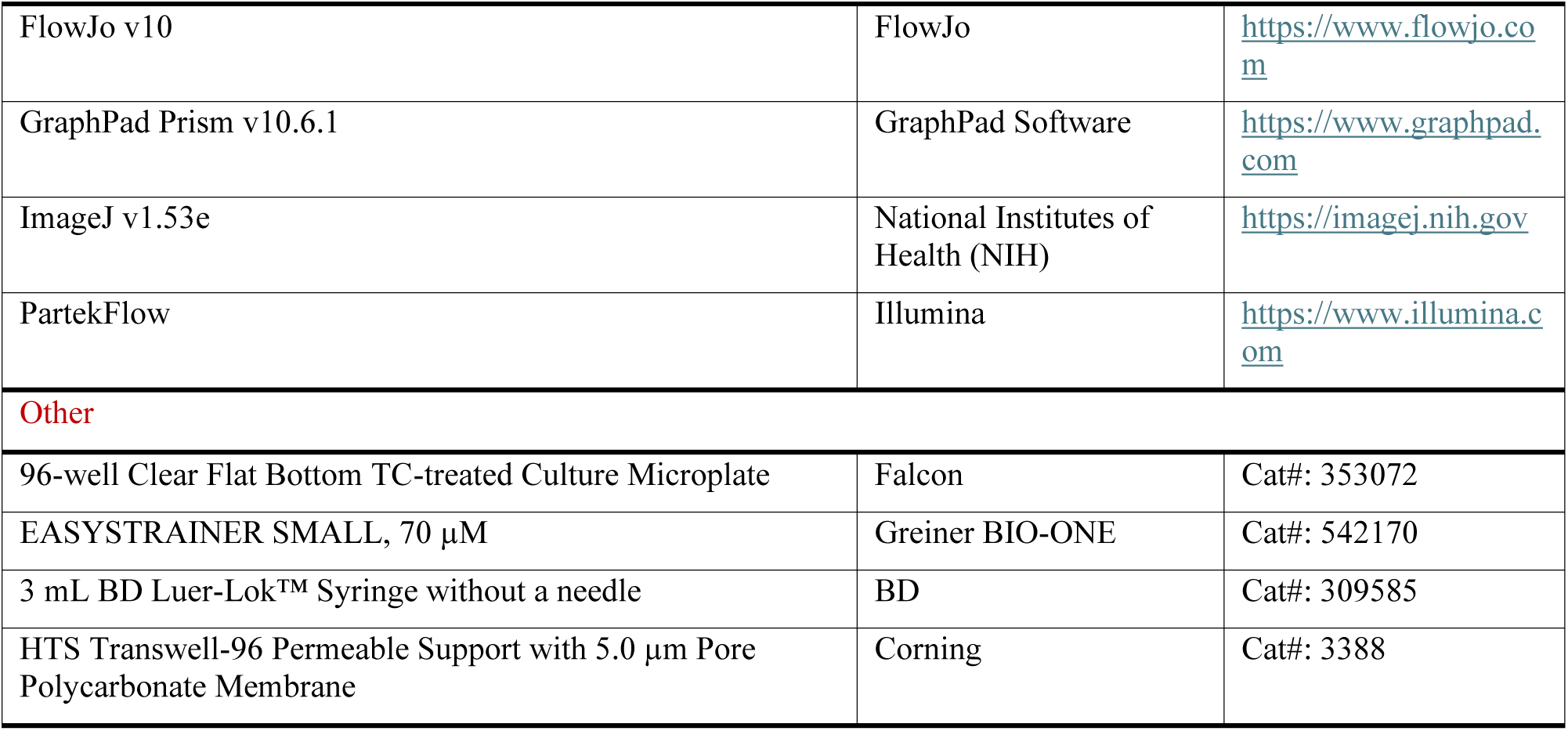

## METHOD DETAILS

### Human blood and tissue samples

All studies involving human samples were approved by the Mayo Clinic Institutional Review Board and conducted in accordance with the Declaration of Helsinki. All samples were also obtained with informed consent (Mayo Clinic-Rochester).

Peripheral blood mononuclear cells (PBMCs) were isolated by density-gradient centrifugation using Ficoll-Paque PLUS (Fisher Scientific) and used either immediately or cryopreserved for subsequent experiments.

Human colon tissue samples from patients undergoing surgery were collected, and macroscopically healthy regions were used for immune cell isolation. Characteristics are listed in **Table S1**. Tonsils were obtained from patients with tonsilitis and were embedded in Tissue-Tek O.C.T. compound (Fisher Scientific) and stored at −80°C until immunofluorescence analysis.

### B cell enrichment and stimulation reagents

CD19^+^ B cells were enriched from PBMCs by negative selection using human pan-B cell enrichment kit (STEMCELL Technologies). Where indicated, IgD^+^ naïve B cells were depleted using human anti-IgD microbeads (Miltenyi Biotec) to obtain untouched CSM B cells. B cells were cultured in complete RPMI 1640 (cRPMI; Gibco) supplemented with 25mM HEPES (Gibco), 1% nonessential amino acids, 2 mM L-glutamate (STEMCELL Technologies), 50 μg/mL penicillin, and 50 μg/mL streptomycin (Gibco), and 10% fetal bovine serum (FBS) (Gemini Bioproducts). For stimulation, anti-BCR (α-IgM + IgA + IgG) at 2.5µg/mL (Jackson ImmunoResearch), IL-2 at 100 ng/mL (PeproTech), IL-15 at 200ng/mL (PeproTech), IL-21 at 20ng/mL (PeproTech), IL-10 at 50ng/mL (PeproTech), IL-27 at 100ng/mL (PeproTech), CpG-B at 2.5µg/mL (ODN 2006, InvivoGen), Poly (I:C) at 10µg/mL (InvivoGen), TLR2-L at 1µg/mL (Pam3CSK4, InvivoGen), and CD40L-L cells^46^ were used.

### Irradiation of CD40L-L cells

CD40L-L cells were seeded at 1 x 10^5^ cells at least 1-2 days before experiments to allow adherence to a flat-bottom 96-well plate. On the day of the experiment, plates were irradiated (RS2000, RadSource) at 50 Gy for 6 min. Cells are then rested for a few hours (2-3h) in the 37°C incubator before adding B cells.

### Flow cytometry and antibodies

Briefly, all cells were first blocked for Fc Receptors (Biolegend) and then stained for surface markers and Live/Dead Fixable Aqua (ThermoFisher) to exclude dead cells. Cells were washed, fixed, and permeabilized, where necessary, using Cytofix/Cytoperm (BD Biosciences) or Foxp3/Transcription Factor Staining Buffer Set (eBioscience) for GVHD flow cytometry assays, followed by intracellular staining for cytokines. For intracellular cytokine detection, cells were stimulated with Phorbol 12-myristate 13-acetate (PMA) 100ng/ml + Ionomycin 1µg/ml (Invivogen) + GolgiPlug containing Brefeldin A and/or Monensin (BD Biosciences) 5 h prior to staining. Samples were acquired using a Cytek Aurora 5L flow cytometer (Cytek Biosciences), and analysis from singlets was performed using FlowJo version v10.10.0 (BD Biosciences). All antibodies used in this study are listed in the **key resources table**.

### Human colon tissue digestion

Single cell suspension of colon tissues was prepared by enzymatic digestion and mechanical disruption. First, colon tissue pieces were incubated in a shaking incubator for 20 min at 37°C in pre-digestion solution containing EDTA (Thermo Scientific) at 5 mM and Dithiothreitol (DTT, Thermo Scientific) at 1 mM in Hank’s Balanced Salt Solution (HBSS, Gibco)^−Ca/−Mg^. This step was repeated twice to remove any mucus and intestinal epithelial cells. Then, pre-digested tissues were further incubated in enzymatic digestion solution containing collagenase D at 500 µg/mL (Roche) and DNase I at 500 µg/mL (Sigma) in HBSS^+Ca/+Mg^ for 1 h at 37°C. Colon pieces were collected in a 70 µM cell strainer on top of a 50 mL falcon tube and mechanically disrupted using 3 mL piston syringe. Cell strainer was rinsed with cRPMI and cells were spun and washed twice with PBS before staining with indicated antibodies.

### Immunofluorescence of human Tonsil

Frozen human tonsils in O.C.T were sectioned at 8 µm thickness. Sections were stained with anti-CD20-FITC (2H7, Biolegend) and anti-TIGIT-PE (A15153G, Biolegend) overnight at 4°C in a humidified chamber. Slides were then counterstained with DAPI (ThermoFisher) and imaged using BZ-X800 microscope (Keyence).

### qRT-PCR

Total RNA was extracted using RNAqueous-Micro Total RNA Isolation Kit (ThermoFisher) according to the manufacturer’s instructions. cDNA was synthesized with iScript Reverse Transcription Supermix for RT-PCR (Biorad) and was amplified using PowerUp SYBR Green Master Mix (ThermoFisher) following the manufacturer’s instructions. qRT-PCR was run on a LightCycler 480 Real-Time PCR System (Roche Diagnostics). RNA expression levels were measured relative to human ribosomal protein S18 (*hRS18*) or the Glyceraldehyde-3-Phosphate Dehydrogenase (*GAPDH)* housekeeping gene. Relative gene expression was calculated using the 2^−ΔΔCt^ method. Primers used are as follows: *hRS18* forward 5’-GCAGAATCCACGCCAGTACAAG-3’ and reverse 5’-GCTTGTTGTCCAGACCATTGGC-3’, *GAPDH* forward 5’-TGTTGCCATCAATGACCCCTT-3’ and reverse 5’-CTCCACGACGTACTCAGCG-3’, *AHR* forward 5’-ACATCACCTACGCCAGTCG-3’ and reverse 5’-CTCTATGCCGCTTGGAAGGAT-3’, *PRDM1* forward 5’-GCTGACAATGATGAATCTCA-3’ and reverse 5’-GTGAAATGTTAGAACGGTAGAG-3’, *IL-10* forward 5’-CCTGGAGGAGGTGATGCCCCA-3’ and reverse 5’-CCTGCTCCACGGCCTTGCTC-3’, *TIGIT* forward 5’-TGGTGGTCATCTGCACAGCAGT-3’ and reverse 5’-TTTCTCCTGAGGTCACCTTCCAC-3’, and *Tim-1* forward 5’-CTTCACCTCAGCCAGCAGAAAC-3’ and reverse 5’-GCCATCTGAAGACTCTGTCACG-3’.

### Generation and purification of TIGIT^+^ CSM B cells

After CD19^+^ B cell enrichment, IgD^+^ naïve B cells were depleted using human anti-IgD microbeads (Miltenyi Biotec) to generate a pool of untouched CSM B cells. CSM B cells were then cultured in the presence of IL-2 and γ-irradiated CD40L-L cells for 6 days. CD19^+^TIGIT^+^ and CD19^+^TIGIT^−^ CSM B cells were sorted by FACS (ARIA III, BD Biosciences).

### Bulk RNA-seq

RNA-seq of purified TBs and CSM B cells was reported previously.^27^ For the comparison between TIGIT ^+^ and TIGIT^−^ CSM B cells, total RNA was isolated from cell lysates using RNAqueous micro total RNA isolation kit (Thermofisher). Total RNA was assessed for quality using the RNA 6000 Pico Kit (Agilent), samples obtained RNA integrity numbers averaging 9.5. Libraries were prepared and sequenced by Novogene using the Illumina NovaSeq PE150 platform. Sequencing reads were aligned to the human reference genome (GRCh38/hg38) using STAR, and gene counts were generated using HTSeq. Genes with counts per million (CPM) <1 were excluded. PCA reduction was performed and differential expression analysis was performed using DESeq2. DEGs were defined as those with an absolute fold change >2 and p < 0.05. GO and KEGG pathway enrichment analyses were performed using significant genes. Data visualization and analysis were performed using Partek Flow software (Illumina).

### Measurement of the effects of STAT5 and PI3K/AKT inhibitors on the induction of TIGIT expression by CSM B cells

Purified CSM B cells were pre-incubated for 1-2h, before culturing them in the presence of γ-irradiated CD40L-L cells and IL-2, with inhibitors for STAT1 at 10µM (Fludarabine, Selleck Chemicals), STAT3 at 20µM (STA-21, Cayman Chemicals), STAT5 at 200µM (AC-4-130, Cayman Chemicals), STAT6 at 100nM (AS1517499, Cayman Chemicals), PI3K at 10µM (LY294002, Cayman Chemicals), Akt at 1µM (MK-2206, Cayman Chemicals), MEK at 10µM (U0126, Cayman Chemicals), and TCF4/β-catenin at 100µM (PNU 74654, Cayman Chemicals). All inhibitors were dissolved in dimethyl sulfoxide (DMSO, Sigma-Aldrich) and final concentration of DMSO was less than 0.005%. Neutralizing α-IL10/IL-10R antibodies at 10µg/mL (Biolegend) were included. The percent inhibition of TIGIT expression was calculated as follows:

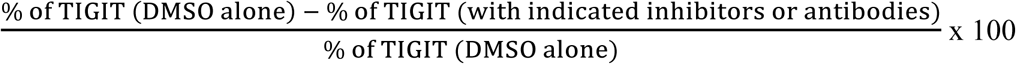

### Stability and Kinetics of TIGIT expression

After 6 days of CSM B cell cultures in the presence of IL-2 and γ-irradiated CD40L-L cells, TIGIT ^+^ and TIGIT^−^ CSM B cells were FACS-sorted and further cultured for additional 6 days in indicated conditions (**Figures 2D-F**). Surface TIGIT expression levels were measured by flow cytometry (Cytek Biosciences). Both frequency of TIGIT^+^ B cells and geometric mean fluorescent intensity (gMFI) of surface TIGIT were assessed.

### TIGIT^+^ CSM B cell-mediated suppression of MDDC activation

Monocytes from the blood of healthy donors were cultured in cRPMI supplemented with GM-CSF (PeproTech) at 100 ng/mL and IL-4 (PeproTech) at 50 ng/mL for 5 days. On days 2 and 4, half of the medium containing the same concentrations of GM-CSF and IL-4 were replenished. FACS-sorted TIGIT ^+^ and TIGIT^−^ CSM B cell populations were co-cultured for two days with MDDCs at 1:2 ratio (B cell : MDDC) in the presence of *E. coli* LPS at 100 ng/mL (InvivoGen). After two days, cells were stimulated for 5 hrs with PMA + ionomycin in the presence of Brefeldin A and then stained for the surface markers CD80, CD83, CD86, CCR7, ICOS-L, and CD40 and intracellular expression of IL-6, TNFα, and IL-12A by flow cytometry. MDDC population was gated based on HLA-DR^+^CD11c^+^. Corresponding antibodies are listed in **key resources table**.

### TIGIT^+^ CSM B cell-mediated suppression of T cell response

Naïve CD4^+^ T cells (purity >95%) from healthy donors were enriched with human naïve CD4^+^ T cell isolation kit (STEMCELL Technologies) and then labeled with CTV (ThermoFisher) for 20 mins in PBS at 37°C. FACS-sorted TIGIT ^+^ and TIGIT^−^ CSM B cell populations were co-cultured with CTV-labeled naïve CD4^+^ T cells at 1:1 ratio (B cell : T cell). CD4^+^ T cells were stimulated with anti-CD3/CD28 dynabeads (ThermoFisher) at 1:1 ratio (cell : bead). After 6 days, CD4^+^ T cell proliferation was assessed by flow cytometry. For intracellular IFN-γ and IL-17A detection, cells were stimulated prior staining for 5 hrs with PMA + ionomycin in the presence of Brefeldin A. The frequency of IFN-γ^+^ and IL-17A^+^ CD4^+^ T cells were measured by flow cytometry. CD4^+^ T population was gated based on CD3^+^CD4^+^. Corresponding antibodies are listed in **key resources table**.

### Anti-TIGIT blocking experiment

In both MDDC and T cell suppression assay, TIGIT^+^ CSM B cells were pre-incubated with α-TIGIT (A15153A, Biolegend) at 10μg/mL or control antibodies for 2 h before co-culturing them with either MDDCs or CTV-labeled naïve CD4^+^ T cells.

### Enzyme-linked immunosorbent assay (ELISA)

FACS-sorted TIGIT ^+^ and TIGIT^−^ CSM B cells (1 x 10^5^ cells/100 µL cRPMI/well) were cultured for three days in 96-well plates in the presence of CD40L-L cells supplemented with IL-2. Culture supernatants were collected, and the amounts of IgM, IgG, IgA, IgA1, IgA2, and total Igs (IgM + IgG + IgA) were quantified by ELISA. Plates were coated with unconjugated α-human IgM, IgG, IgA, IgA1 or IgA2 capture antibodies (SouthernBiotech), followed by detection with HRP-conjugated secondary antibodies (SouthernBiotech). TMB substrate (BD Biosciences) was used for signal development. To determine the Ig titer, culture supernatants were analyzed undiluted and in 2-fold serial dilutions (1:2 to 1:256). Endpoint titers were defined as the highest dilution yielding an optical density (OD) above a predetermined cutoff, calculated as the mean OD of blank wells plus two standard deviations. Titers of individual Ig isotypes were subsequently normalized to the total Ig titer.

### Chemotaxis assay

FACS-sorted TIGIT ^+^ and TIGIT^−^ CSM B cells (2 x 10^4^ cells/75 µL cRPMI/well) were added in the upper chamber of 96-transwell plates (Corning) containing 235 µL cRPMI of CCL25 chemokine (PeproTech) at indicated concentrations in the lower chamber. After 2 h of incubation at 37°C, to allow migration through the 5 µm pore-size membrane, cells in the lower chamber were harvested and counted by flow cytometry (Cytek Biosciences) using CountBright Plus Absolute counting beads (ThermoFisher).

### aGVHD and TIGIT^+^ CSM B cell treatment

Female NOG mice (NOD.Cg-*Prkdc^scid^ Il2rg^tm1Sug^*/JicTac; Taconic Biosciences) aged 8 to 10 weeks were used. All animals received indicated numbers of freshly isolated PBMCs from healthy adult individuals. Animals were treated with indicated subsets of B cells (TIGIT^+^ or TIGIT^−^ CSM B cells) via i.v. in tail vein. Mice were monitored for assessing the GVHD scores at least three times per week. Clinical symptoms of aGVHD were graded according to published criteria.^79^ Peripheral blood was collected weekly after PBMC transplantation. Mice were euthanized if weight loss exceeded 25% of initial body weight or if the cumulative clinical score reached >5-6. At sacrifice, spleen and tissues, including liver and intestines, were collected for flow cytometry and histological analysis. Splenocytes were stimulated prior staining for 5 hrs with PMA + ionomycin in the presence of Brefeldin A and Monensin. Antibodies are listed in **key resources table**.

All procedures were performed under an approved protocol and in accordance with the Mayo Clinic Institutional Animal Care and Use Committee guidelines for proper care and use of laboratory animals.

### Statistical analysis

Data are presented as mean ± SEM unless otherwise indicated. Statistical analyses were performed using GraphPad Prism (version v10.10.0). Comparisons between two groups were performed using two-tailed unpaired or paired Student’s t-tests, whereas comparisons among multiple groups were performed using one-way or two-way ANOVA with appropriate post hoc testing. Statistical significance for Kaplan–Meier survival curves was determined using the log-rank (Mantel–Cox) test. P-values are represented by the following scale: ns, not significant > 0.05, ∗ <0.05, ∗∗ <0.01, ∗∗∗ <0.001, ∗∗∗∗ <0.0001.

## REFERENCES

1. Blair, P.A., Norena, L.Y., Flores-Borja, F., Rawlings, D.J., Isenberg, D.A., Ehrenstein, M.R., and Mauri, C. (2010). CD19(+)CD24(hi)CD38(hi) B cells exhibit regulatory capacity in healthy individuals but are functionally impaired in systemic Lupus Erythematosus patients. Immunity 32, 129–140. 10.1016/j.immuni.2009.11.009.

2. Mauri, C., and Menon, M. (2017). Human regulatory B cells in health and disease: therapeutic potential. J Clin Invest 127, 772–779. 10.1172/JCI85113.

3. Khoder, A., Sarvaria, A., Alsuliman, A., Chew, C., Sekine, T., Cooper, N., Mielke, S., de Lavallade, H., Muftuoglu, M., Fernandez Curbelo, I., et al. (2014). Regulatory B cells are enriched within the IgM memory and transitional subsets in healthy donors but are deficient in chronic GVHD. Blood 124, 2034–2045. 10.1182/blood-2014-04-571125.

4. Yanaba, K., Yoshizaki, A., Asano, Y., Kadono, T., Tedder, T.F., and Sato, S. (2011). IL-10-producing regulatory B10 cells inhibit intestinal injury in a mouse model. Am J Pathol 178, 735–743. 10.1016/j.ajpath.2010.10.022.

5. DiLillo, D.J., Matsushita, T., and Tedder, T.F. (2010). B10 cells and regulatory B cells balance immune responses during inflammation, autoimmunity, and cancer. Ann N Y Acad Sci 1183, 38–57. 10.1111/j.1749-6632.2009.05137.x.

6. Asashima, H., Axisa, P.P., Pham, T.H.G., Longbrake, E.E., Ruff, W.E., Lele, N., Cohen, I., Raddassi, K., Sumida, T.S., and Hafler, D.A. (2022). Impaired TIGIT expression on B cells drives circulating follicular helper T cell expansion in multiple sclerosis. J Clin Invest 132. 10.1172/JCI156254.

7. Bottomley, M.J., Brook, M.O., Shankar, S., Hester, J., and Issa, F. (2022). Towards regulatory cellular therapies in solid organ transplantation. Trends Immunol 43, 8–21. 10.1016/j.it.2021.11.001.

8. Slepicka, P.F., Yazdanifar, M., and Bertaina, A. (2021). Harnessing Mechanisms of Immune Tolerance to Improve Outcomes in Solid Organ Transplantation: A Review. Front Immunol 12, 688460. 10.3389/fimmu.2021.688460.

9. Baert, L., Mahmudul, H.M., Stegall, M., Joo, H., and Oh, S. (2024). B Cell-mediated Immune Regulation and the Quest for Transplantation Tolerance. Transplantation 108, 2021–2033. 10.1097/TP.0000000000004948.

10. Wortel, C.M., and Heidt, S. (2017). Regulatory B cells: Phenotype, function and role in transplantation. Transpl Immunol 41, 1–9. 10.1016/j.trim.2017.02.004.

11. Cherukuri, A., Mohib, K., and Rothstein, D.M. (2021). Regulatory B cells: TIM-1, transplant tolerance, and rejection. Immunol Rev 299, 31–44. 10.1111/imr.12933.

12. Shen, P., Roch, T., Lampropoulou, V., O’Connor, R.A., Stervbo, U., Hilgenberg, E., Ries, S., Dang, V.D., Jaimes, Y., Daridon, C., et al. (2014). IL-35-producing B cells are critical regulators of immunity during autoimmune and infectious diseases. Nature 507, 366–370. 10.1038/nature12979.

13. Gong, Y., Zhao, C., Zhao, P., Wang, M., Zhou, G., Han, F., Cui, Y., Qian, J., Zhang, H., Xiong, H., et al. (2015). Role of IL-10-Producing Regulatory B Cells in Chronic Hepatitis B Virus Infection. Dig Dis Sci 60, 1308–1314. 10.1007/s10620-014-3358-1.

14. Liu, Y., Cheng, L.S., Wu, S.D., Wang, S.Q., Li, L., She, W.M., Li, J., Wang, J.Y., and Jiang, W. (2016). IL-10-producing regulatory B-cells suppressed effector T-cells but enhanced regulatory T-cells in chronic HBV infection. Clin Sci (Lond) 130, 907–919. 10.1042/CS20160069.

15. Liu, Y., Chen, Y., Li, Z., Han, Y., Sun, Y., Wang, Q., Liu, B., and Su, Z. (2013). Role of IL-10-producing regulatory B cells in control of cerebral malaria in Plasmodium berghei infected mice. Eur J Immunol 43, 2907–2918. 10.1002/eji.201343512.

16. Das, A., Ellis, G., Pallant, C., Lopes, A.R., Khanna, P., Peppa, D., Chen, A., Blair, P., Dusheiko, G., Gill, U., et al. (2012). IL-10-producing regulatory B cells in the pathogenesis of chronic hepatitis B virus infection. J Immunol 189, 3925–3935. 10.4049/jimmunol.1103139.

17. Horikawa, M., Minard-Colin, V., Matsushita, T., and Tedder, T.F. (2011). Regulatory B cell production of IL-10 inhibits lymphoma depletion during CD20 immunotherapy in mice. J Clin Invest 121, 4268–4280. 10.1172/JCI59266.

18. Shao, Y., Lo, C.M., Ling, C.C., Liu, X.B., Ng, K.T., Chu, A.C., Ma, Y.Y., Li, C.X., Fan, S.T., and Man, K. (2014). Regulatory B cells accelerate hepatocellular carcinoma progression via CD40/CD154 signaling pathway. Cancer Lett 355, 264–272. 10.1016/j.canlet.2014.09.026.

19. Olkhanud, P.B., Damdinsuren, B., Bodogai, M., Gress, R.E., Sen, R., Wejksza, K., Malchinkhuu, E., Wersto, R.P., and Biragyn, A. (2011). Tumor-evoked regulatory B cells promote breast cancer metastasis by converting resting CD4(+) T cells to T-regulatory cells. Cancer Res 71, 3505–3515. 10.1158/0008-5472.CAN-10-4316.

20. Schioppa, T., Moore, R., Thompson, R.G., Rosser, E.C., Kulbe, H., Nedospasov, S., Mauri, C., Coussens, L.M., and Balkwill, F.R. (2011). B regulatory cells and the tumor-promoting actions of TNF-alpha during squamous carcinogenesis. Proc Natl Acad Sci U S A 108, 10662–10667. 10.1073/pnas.1100994108.

21. Catalan, D., Mansilla, M.A., Ferrier, A., Soto, L., Oleinika, K., Aguillon, J.C., and Aravena, O. (2021). Immunosuppressive Mechanisms of Regulatory B Cells. Front Immunol 12, 611795. 10.3389/fimmu.2021.611795.

22. Nouel, A., Pochard, P., Simon, Q., Segalen, I., Le Meur, Y., Pers, J.O., and Hillion, S. (2015). B-Cells induce regulatory T cells through TGF-beta/IDO production in A CTLA-4 dependent manner. J Autoimmun 59, 53–60. 10.1016/j.jaut.2015.02.004.

23. Figueiro, F., Muller, L., Funk, S., Jackson, E.K., Battastini, A.M., and Whiteside, T.L. (2016). Phenotypic and functional characteristics of CD39(high) human regulatory B cells (Breg). Oncoimmunology 5, e1082703. 10.1080/2162402X.2015.1082703.

24. Kaku, H., Cheng, K.F., Al-Abed, Y., and Rothstein, T.L. (2014). A novel mechanism of B cell-mediated immune suppression through CD73 expression and adenosine production. J Immunol 193, 5904–5913. 10.4049/jimmunol.1400336.

25. Choi, J.K., Yu, C.R., Bing, S.J., Jittayasothorn, Y., Mattapallil, M.J., Kang, M., Park, S.B., Lee, H.S., Dong, L., Shi, G., et al. (2021). IL-27-producing B-1a cells suppress neuroinflammation and CNS autoimmune diseases. Proc Natl Acad Sci U S A 118. 10.1073/pnas.2109548118.

26. Goode, I., Xu, H., and Ildstad, S.T. (2014). Regulatory B Cells: The New “It” Cell. Transpl P 46, 3–8. 10.1016/j.transproceed.2013.08.075.

27. Hasan, M.M., Nair, S.S., O’Leary, J.G., Thompson-Snipes, L., Nyarige, V., Wang, J., Park, W., Stegall, M., Heilman, R., Klintmalm, G.B., et al. (2021). Implication of TIGIT(+) human memory B cells in immune regulation. Nat Commun 12, 1534. 10.1038/s41467-021-21413-y.

28. Xiao, S., Bod, L., Pochet, N., Kota, S.B., Hu, D., Madi, A., Kilpatrick, J., Shi, J., Ho, A., Zhang, H., et al. (2020). Checkpoint Receptor TIGIT Expressed on Tim-1(+) B Cells Regulates Tissue Inflammation. Cell Rep 32, 107892. 10.1016/j.celrep.2020.107892.

29. Anderson, A.C., Joller, N., and Kuchroo, V.K. (2016). Lag-3, Tim-3, and TIGIT: Co-inhibitory Receptors with Specialized Functions in Immune Regulation. Immunity 44, 989–1004. 10.1016/j.immuni.2016.05.001.

30. Joller, N., Hafler, J.P., Brynedal, B., Kassam, N., Spoerl, S., Levin, S.D., Sharpe, A.H., and Kuchroo, V.K. (2011). Cutting edge: TIGIT has T cell-intrinsic inhibitory functions. J Immunol 186, 1338–1342. 10.4049/jimmunol.1003081.

31. Joller, N., Lozano, E., Burkett, P.R., Patel, B., Xiao, S., Zhu, C., Xia, J., Tan, T.G., Sefik, E., Yajnik, V., et al. (2014). Treg cells expressing the coinhibitory molecule TIGIT selectively inhibit proinflammatory Th1 and Th17 cell responses. Immunity 40, 569–581. 10.1016/j.immuni.2014.02.012.

32. Lozano, E., Dominguez-Villar, M., Kuchroo, V., and Hafler, D.A. (2012). The TIGIT/CD226 axis regulates human T cell function. J Immunol 188, 3869–3875. 10.4049/jimmunol.1103627.

33. Yu, X., Harden, K., Gonzalez, L.C., Francesco, M., Chiang, E., Irving, B., Tom, I., Ivelja, S., Refino, C.J., Clark, H., et al. (2009). The surface protein TIGIT suppresses T cell activation by promoting the generation of mature immunoregulatory dendritic cells. Nat Immunol 10, 48–57. 10.1038/ni.1674.

34. Varghese, J.F., Kaskow, B.J., von Glehn, F., Case, J., Li, Z., Jule, A.M., Berdan, E., Ho Sui, S.J., Hu, Y., Krishnan, R., et al. (2024). Human regulatory memory B cells defined by expression of TIM-1 and TIGIT are dysfunctional in multiple sclerosis. Front Immunol 15, 1360219. 10.3389/fimmu.2024.1360219.

35. DeLong, J.H., O’Hara Hall, A., Rausch, M., Moodley, D., Perry, J., Park, J., Phan, A.T., Beiting, D.P., Kedl, R.M., Hill, J.A., and Hunter, C.A. (2019). IL-27 and TCR Stimulation Promote T Cell Expression of Multiple Inhibitory Receptors. Immunohorizons 3, 13–25. 10.4049/immunohorizons.1800083.

36. Morrow, K.N., Liang, Z., Xue, M., Chihade, D.B., Sun, Y., Chen, C.W., Coopersmith, C.M., and Ford, M.L. (2021). The IL-27 receptor regulates TIGIT on memory CD4(+) T cells during sepsis. iScience 24, 102093. 10.1016/j.isci.2021.102093.

37. Zhang, H., Madi, A., Yosef, N., Chihara, N., Awasthi, A., Pot, C., Lambden, C., Srivastava, A., Burkett, P.R., Nyman, J., et al. (2020). An IL-27-Driven Transcriptional Network Identifies Regulators of IL-10 Expression across T Helper Cell Subsets. Cell Rep 33, 108433. 10.1016/j.celrep.2020.108433.

38. Chihara, N., Madi, A., Kondo, T., Zhang, H., Acharya, N., Singer, M., Nyman, J., Marjanovic, N.D., Kowalczyk, M.S., Wang, C., et al. (2018). Induction and transcriptional regulation of the co-inhibitory gene module in T cells. Nature 558, 454–459. 10.1038/s41586-018-0206-z.

39. Komegae, E.N., Grund, L.Z., Lopes-Ferreira, M., and Lima, C. (2013). TLR2, TLR4 and the MyD88 signaling are crucial for the in vivo generation and the longevity of long-lived antibody-secreting cells. PLoS One 8, e71185. 10.1371/journal.pone.0071185.

40. Marshall-Clarke, S., Downes, J.E., Haga, I.R., Bowie, A.G., Borrow, P., Pennock, J.L., Grencis, R.K., and Rothwell, P. (2007). Polyinosinic acid is a ligand for toll-like receptor 3. J Biol Chem 282, 24759–24766. 10.1074/jbc.M700188200.

41. Sackesen, C., van de Veen, W., Akdis, M., Soyer, O., Zumkehr, J., Ruckert, B., Stanic, B., Kalayci, O., Alkan, S.S., Gursel, I., and Akdis, C.A. (2013). Suppression of B-cell activation and IgE, IgA, IgG1 and IgG4 production by mammalian telomeric oligonucleotides. Allergy 68, 593–603. 10.1111/all.12133.

42. Bernasconi, N.L., Onai, N., and Lanzavecchia, A. (2003). A role for Toll-like receptors in acquired immunity: up-regulation of TLR9 by BCR triggering in naive B cells and constitutive expression in memory B cells. Blood 101, 4500–4504. 10.1182/blood-2002-11-3569.

43. Jones, D.M., Read, K.A., and Oestreich, K.J. (2020). Dynamic Roles for IL-2-STAT5 Signaling in Effector and Regulatory CD4(+) T Cell Populations. J Immunol 205, 1721–1730. 10.4049/jimmunol.2000612.

44. Gauthier, J., Maugendre, M., Leonard, S., Desvois, Y., Pineau, M., Pinon, R., Hipp, N., Tarte, K., Ame, P., Hillion, S., et al. (2026). B cell-intrinsic IL-2 signaling regulates inflammation by promoting IL-10 expression in CD25(+) age-associated B cells. Immunity 59, 354–371 e359. 10.1016/j.immuni.2026.01.015.

45. Elgueta, R., Benson, M.J., de Vries, V.C., Wasiuk, A., Guo, Y., and Noelle, R.J. (2009). Molecular mechanism and function of CD40/CD40L engagement in the immune system. Immunol Rev 229, 152–172. 10.1111/j.1600-065X.2009.00782.x.

46. Arpin, C., Dechanet, J., Van Kooten, C., Merville, P., Grouard, G., Briere, F., Banchereau, J., and Liu, Y.J. (1995). Generation of memory B cells and plasma cells in vitro. Science 268, 720–722. 10.1126/science.7537388.

47. Piper, C.J.M., Rosser, E.C., Oleinika, K., Nistala, K., Krausgruber, T., Rendeiro, A.F., Banos, A., Drozdov, I., Villa, M., Thomson, S., et al. (2019). Aryl Hydrocarbon Receptor Contributes to the Transcriptional Program of IL-10-Producing Regulatory B Cells. Cell Rep 29, 1878–1892 e1877. 10.1016/j.celrep.2019.10.018.

48. Johansson-Lindbom, B., and Agace, W.W. (2007). Generation of gut-homing T cells and their localization to the small intestinal mucosa. Immunol Rev 215, 226–242. 10.1111/j.1600-065X.2006.00482.x.

49. Jarmin, D.I., Rits, M., Bota, D., Gerard, N.P., Graham, G.J., Clark-Lewis, I., and Gerard, C. (2000). Cutting edge: identification of the orphan receptor G-protein-coupled receptor 2 as CCR10, a specific receptor for the chemokine ESkine. J Immunol 164, 3460–3464. 10.4049/jimmunol.164.7.3460.

50. Matloubian, M., Lo, C.G., Cinamon, G., Lesneski, M.J., Xu, Y., Brinkmann, V., Allende, M.L., Proia, R.L., and Cyster, J.G. (2004). Lymphocyte egress from thymus and peripheral lymphoid organs is dependent on S1P receptor 1. Nature 427, 355–360. 10.1038/nature02284.

51. Cyster, J.G., and Schwab, S.R. (2012). Sphingosine-1-phosphate and lymphocyte egress from lymphoid organs. Annu Rev Immunol 30, 69–94. 10.1146/annurev-immunol-020711-075011.

52. Sallusto, F., Lenig, D., Forster, R., Lipp, M., and Lanzavecchia, A. (1999). Two subsets of memory T lymphocytes with distinct homing potentials and effector functions. Nature 401, 708–712. 10.1038/44385.

53. Bonecchi, R., Bianchi, G., Bordignon, P.P., D’Ambrosio, D., Lang, R., Borsatti, A., Sozzani, S., Allavena, P., Gray, P.A., Mantovani, A., and Sinigaglia, F. (1998). Differential expression of chemokine receptors and chemotactic responsiveness of type 1 T helper cells (Th1s) and Th2s. J Exp Med 187, 129–134. 10.1084/jem.187.1.129.

54. Nakayama, T., Hieshima, K., Nagakubo, D., Sato, E., Nakayama, M., Kawa, K., and Yoshie, O. (2004). Selective induction of Th2-attracting chemokines CCL17 and CCL22 in human B cells by latent membrane protein 1 of Epstein-Barr virus. J Virol 78, 1665–1674. 10.1128/jvi.78.4.1665-1674.2004.

55. Morgan, D.A., Ruscetti, F.W., and Gallo, R. (1976). Selective in vitro growth of T lymphocytes from normal human bone marrows. Science 193, 1007–1008. 10.1126/science.181845.

56. Robb, R.J., Munck, A., and Smith, K.A. (1981). T cell growth factor receptors. Quantitation, specificity, and biological relevance. J Exp Med 154, 1455–1474. 10.1084/jem.154.5.1455.

57. Rosenberg, S.A. (2014). IL-2: the first effective immunotherapy for human cancer. J Immunol 192, 5451–5458. 10.4049/jimmunol.1490019.

58. Malek, T.R., Yu, A., Vincek, V., Scibelli, P., and Kong, L. (2002). CD4 regulatory T cells prevent lethal autoimmunity in IL-2Rbeta-deficient mice. Implications for the nonredundant function of IL-2. Immunity 17, 167–178. 10.1016/s1074-7613(02)00367-9.

59. Spolski, R., Li, P., and Leonard, W.J. (2018). Biology and regulation of IL-2: from molecular mechanisms to human therapy. Nat Rev Immunol 18, 648–659. 10.1038/s41577-018-0046-y.

60. Chinen, T., Kannan, A.K., Levine, A.G., Fan, X., Klein, U., Zheng, Y., Gasteiger, G., Feng, Y., Fontenot, J.D., and Rudensky, A.Y. (2016). An essential role for the IL-2 receptor in T(reg) cell function. Nat Immunol 17, 1322–1333. 10.1038/ni.3540.

61. Josefowicz, S.Z., Lu, L.F., and Rudensky, A.Y. (2012). Regulatory T cells: mechanisms of differentiation and function. Annu Rev Immunol 30, 531–564. 10.1146/annurev.immunol.25.022106.141623.

62. Majri, S.S., Fritz, J.M., Villarino, A.V., Zheng, L., Kanellopoulou, C., Chaigne-Delalande, B., Gronholm, J., Niemela, J.E., Afzali, B., Biancalana, M., et al. (2018). STAT5B: A Differential Regulator of the Life and Death of CD4(+) Effector Memory T Cells. J Immunol 200, 110–118. 10.4049/jimmunol.1701133.

63. Bishop, G.A., and Hostager, B.S. (2001). Signaling by CD40 and its mimics in B cell activation. Immunol Res 24, 97–109. 10.1385/IR:24:2:097.

64. Shin, M., Yi, E.H., Kim, B.H., Shin, J.C., Park, J.Y., Cho, C.H., Park, J.W., Choi, K.Y., and Ye, S.K. (2016). STAT3 Potentiates SIAH-1 Mediated Proteasomal Degradation of beta-Catenin in Human Embryonic Kidney Cells. Mol Cells 39, 821–826. 10.14348/molcells.2016.0212.

65. Schmetterer, K.G., Neunkirchner, A., and Pickl, W.F. (2012). Naturally occurring regulatory T cells: markers, mechanisms, and manipulation. FASEB J 26, 2253–2276. 10.1096/fj.11-193672.

66. Shevach, E.M. (2009). Mechanisms of foxp3+ T regulatory cell-mediated suppression. Immunity 30, 636–645. 10.1016/j.immuni.2009.04.010.

67. Read, S., Malmstrom, V., and Powrie, F. (2000). Cytotoxic T lymphocyte-associated antigen 4 plays an essential role in the function of CD25(+)CD4(+) regulatory cells that control intestinal inflammation. J Exp Med 192, 295–302. 10.1084/jem.192.2.295.

68. Iwata, Y., Matsushita, T., Horikawa, M., Dilillo, D.J., Yanaba, K., Venturi, G.M., Szabolcs, P.M., Bernstein, S.H., Magro, C.M., Williams, A.D., et al. (2011). Characterization of a rare IL-10-competent B-cell subset in humans that parallels mouse regulatory B10 cells. Blood 117, 530–541. 10.1182/blood-2010-07-294249.

69. Mauri, C., and Menon, M. (2015). The expanding family of regulatory B cells. Int Immunol 27, 479–486. 10.1093/intimm/dxv038.

70. Woof, J.M., and Kerr, M.A. (2006). The function of immunoglobulin A in immunity. J Pathol 208, 270–282. 10.1002/path.1877.

71. Woof, J.M., and Russell, M.W. (2011). Structure and function relationships in IgA. Mucosal Immunol 4, 590–597. 10.1038/mi.2011.39.

72. Cerutti, A. (2008). The regulation of IgA class switching. Nat Rev Immunol 8, 421–434. 10.1038/nri2322.

73. Rosser, E.C., Oleinika, K., Tonon, S., Doyle, R., Bosma, A., Carter, N.A., Harris, K.A., Jones, S.A., Klein, N., and Mauri, C. (2014). Regulatory B cells are induced by gut microbiota-driven interleukin-1beta and interleukin-6 production. Nat Med 20, 1334–1339. 10.1038/nm.3680.

74. Cerutti, A., Cols, M., and Puga, I. (2013). Marginal zone B cells: virtues of innate-like antibody-producing lymphocytes. Nat Rev Immunol 13, 118–132. 10.1038/nri3383.

75. Shalapour, S., Font-Burgada, J., Di Caro, G., Zhong, Z., Sanchez-Lopez, E., Dhar, D., Willimsky, G., Ammirante, M., Strasner, A., Hansel, D.E., et al. (2015). Immunosuppressive plasma cells impede T-cell-dependent immunogenic chemotherapy. Nature 521, 94–98. 10.1038/nature14395.

76. Rosser, E.C., and Mauri, C. (2015). Regulatory B cells: origin, phenotype, and function. Immunity 42, 607–612. 10.1016/j.immuni.2015.04.005.

77. Menon, M., Blair, P.A., Isenberg, D.A., and Mauri, C. (2016). A Regulatory Feedback between Plasmacytoid Dendritic Cells and Regulatory B Cells Is Aberrant in Systemic Lupus Erythematosus. Immunity 44, 683–697. 10.1016/j.immuni.2016.02.012.

78. Mizoguchi, A., and Bhan, A.K. (2006). A case for regulatory B cells. J Immunol 176, 705–710. 10.4049/jimmunol.176.2.705.

79. Cooke, K.R., Kobzik, L., Martin, T.R., Brewer, J., Delmonte, J., Jr., Crawford, J.M., and Ferrara, J.L. (1996). An experimental model of idiopathic pneumonia syndrome after bone marrow transplantation: I. The roles of minor H antigens and endotoxin. Blood 88, 3230–3239.

