## Supplemental information for "IL-2 and CD40L can establish and sustain a human B cell-mediated immune regulatory network"

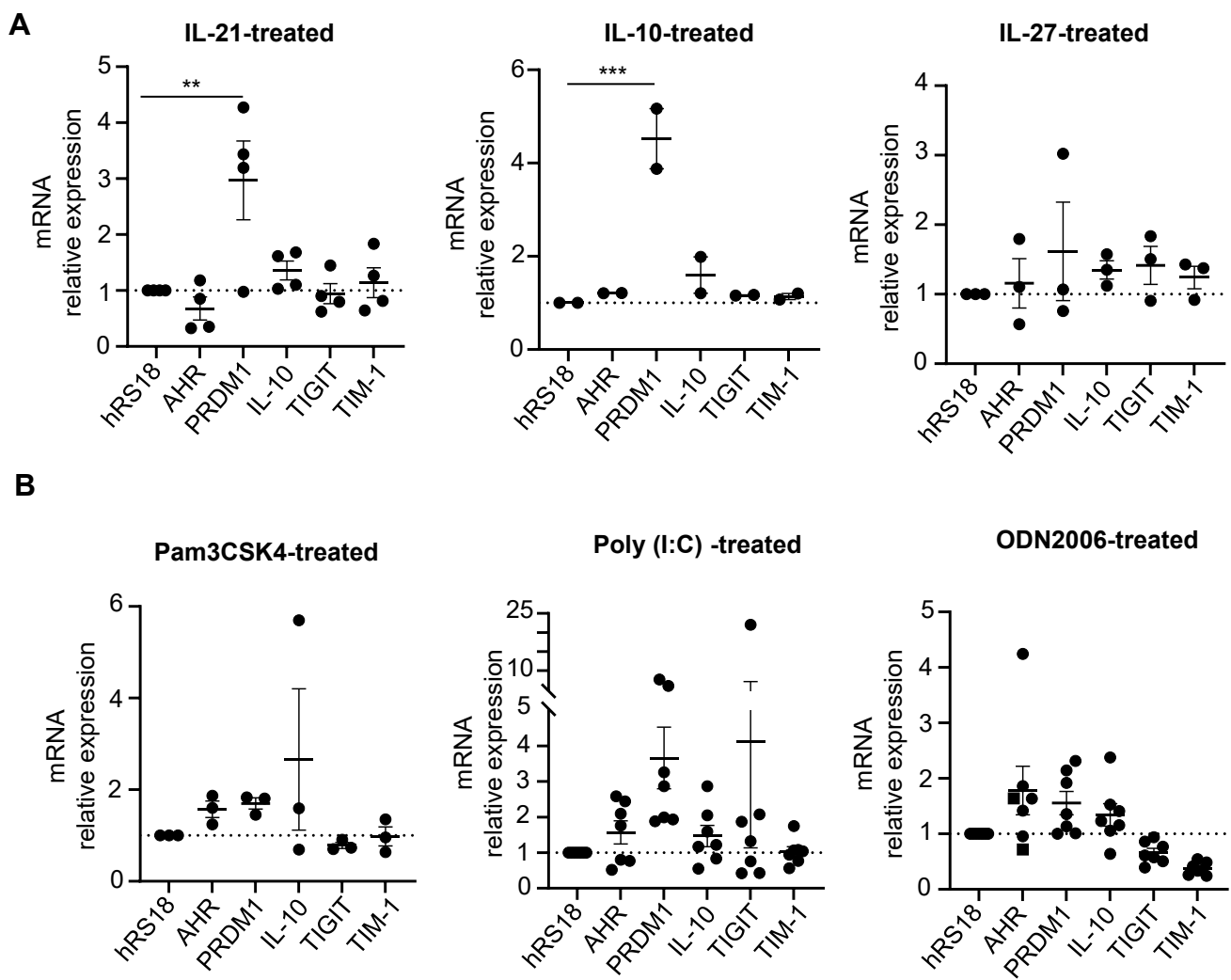

**Figure S1. IL-21, IL-10, IL-27, or ligands of TLRs 2, 3, and 9 do not induce *TIGIT* mRNA expression in B cells. Data associated with Figure 1.**

(A) Purified CD19<sup>+</sup> B cells from the blood of healthy donors were stimulated with α-BCR antibody (2.5 μg/ml) and cultured for 36 hours in the presence of IL-21 (20 ng/ml), IL-10 (50 ng/ml), or IL-27 (100 ng/ml). mRNA expression of *AHR*, *PRDM1*, *IL-10*, *TIGIT*, and *TIM-1* was measured by qPCR. B cells from 2-5 healthy donors were tested. (B) α-BCR-stimulated CD19<sup>+</sup> B cells from the blood of healthy donors were cultured for 36 hours in the presence of the TLR9 ligand ODN2006 (2.5 μg/ml), the TLR2 ligand Pam3CSK4 (1 μg/ml), and the TLR3 ligand poly (I:C) (10 μg/ml). mRNA expression of *AHR*, *PRDM1*, *IL-10*, *TIGIT*, and *TIM-1* was measured by qPCR. B cells from 3-7 healthy donors were tested. Error bars indicate SEM. Statistical significance was determined using one-way ANOVA with Dunnett's multiple comparisons test. \*\* < 0.01, \*\*\* < 0.001. Unless indicated, there was no significant difference.

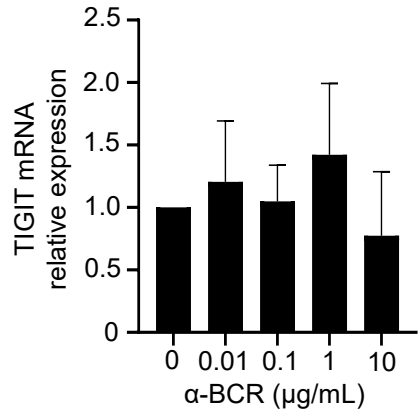

**Figure S2. α-BCR does not significantly alter TIGIT mRNA expression in CD19<sup>+</sup> B cells. Data associated with Figure 1.**

Purified CD19<sup>+</sup> B cells from the blood of 3 healthy donors were stimulated with indicated amount of α-BCR antibodies. After 36 hours, *TIGIT* mRNA expression level was quantified by qPCR. Error bars indicate SEM of triplicate assay. Statistical significance was determined using one-way ANOVA with Dunnett's multiple comparisons test. All comparisons are not significant.

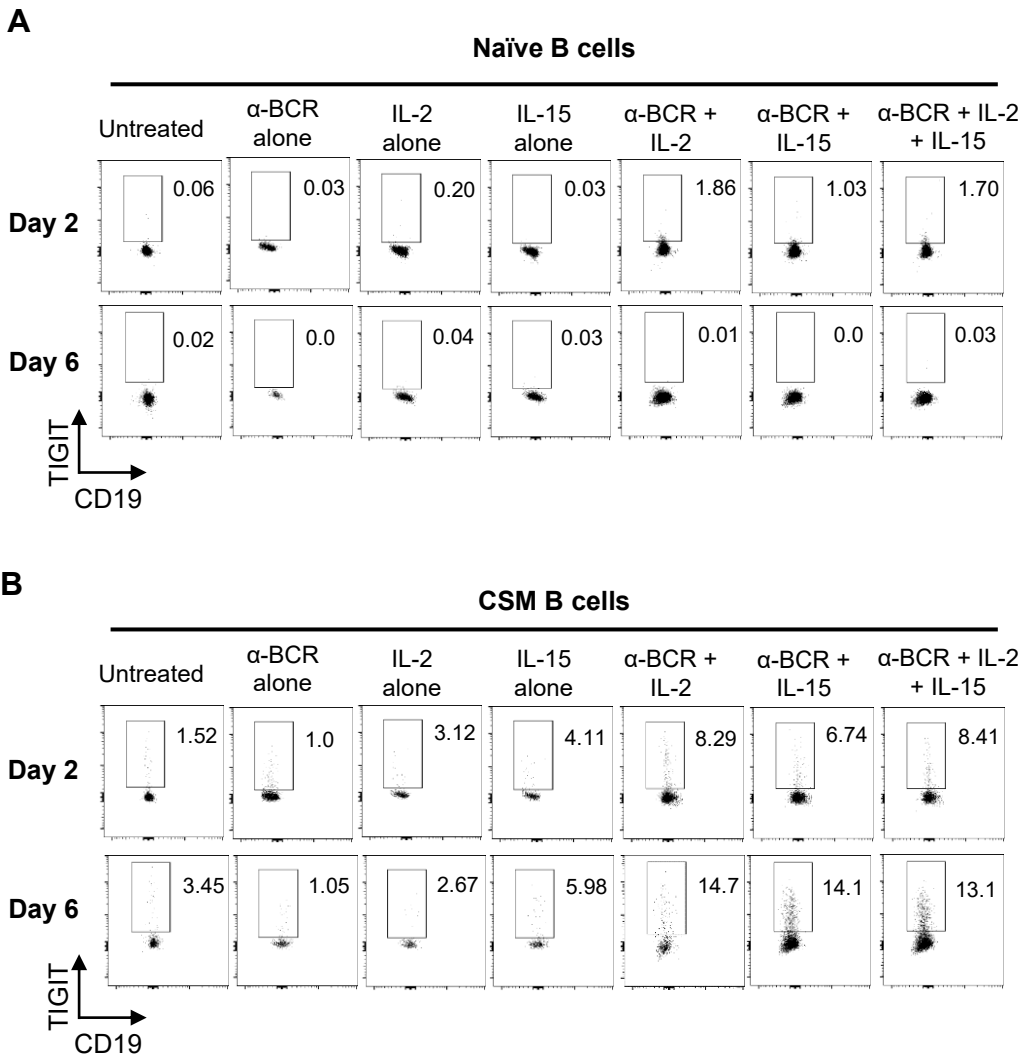

**Figure S3. IL-2 and IL-15 promote surface TIGIT expression on α-BCR-activated CSM B cells. Data associated with Figure 1.**

(A) Purified naïve and (B) CSM B cells were cultured for 2 and 6 days under the indicated conditions. Representative FACS plots are presented. Treatment with α-BCR, IL-2 and IL-15 alone were added as additional controls.

**A**

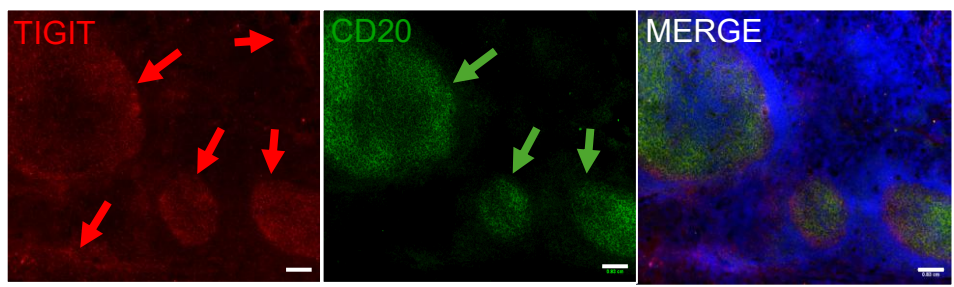

**B**

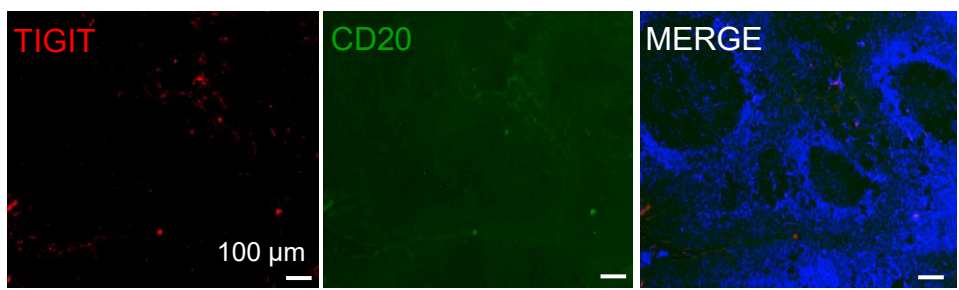

**Figure S4. Immunofluorescence staining of inflamed human tonsil tissue sections with anti-TIGIT, anti-CD20, and DAPI. Data associated with Figure 1.**

(A) Tonsil section stained with anti-TIGIT, anti-CD20, and DAPI.

(B) Tonsil section stained with control antibodies and DAPI.

Magnification x4. Bar size 100 μm. Arrows indicate the major staining patterns described in the text

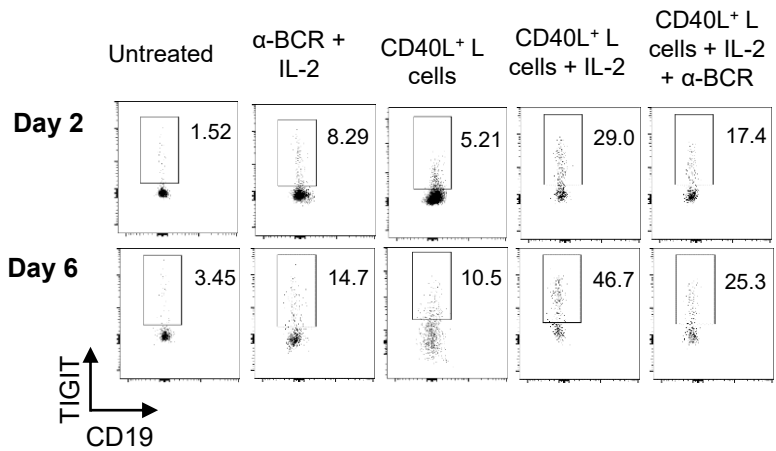

**Figure S5. CD40L further promotes surface TIGIT expression on CSM B cells. Data associated with Figure 2.**

Purified CSM B cells were cultured for 2 and 6 days under the indicated conditions. Representative FACS plots are presented.

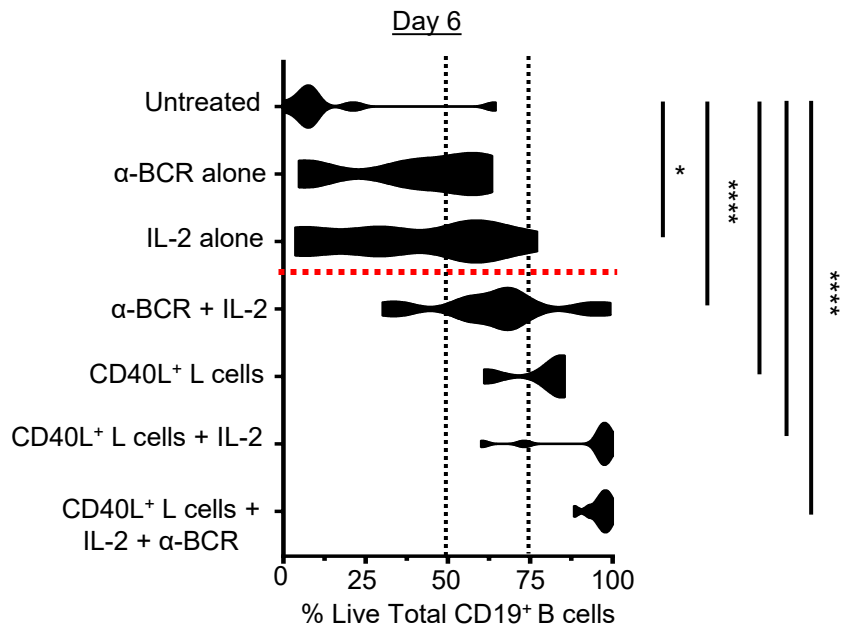

**Figure S6. CD40L support survival of TIGIT<sup>+</sup> B cells. Data associated with Figure 2.**

Percentage of live cells from total CD19<sup>+</sup> B cells was measured by flow cytometry at day 6 with indicated stimuli. Data are presented as min–max values with individual donor data points. Statistical analyses were performed using one-way ANOVA with Dunnett's multiple-comparisons test. Unless indicated, there was no significant difference.

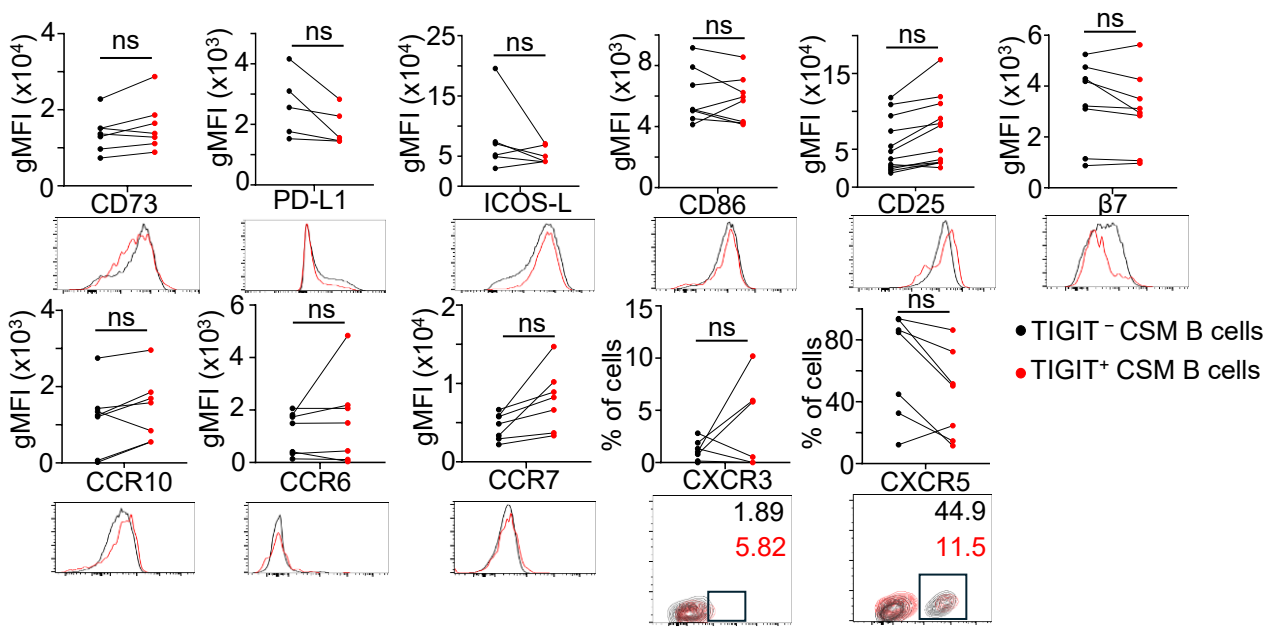

**Figure S7. Cell surface expression of inhibitory/activation markers and chemokine receptors on TIGIT<sup>+</sup> and TIGIT<sup>-</sup> CSM B cells. Data associated with Figure 4.**

Enriched pool of untouched CSM B cells were stimulated with CD40L<sup>+</sup> L cells + IL-2 for 6 days. Cell surface expression of CD73, PD-L1, ICOSL, CD86, CD25, β7 integrin, CCR10, CCR6, CCR7, CXCR3, and CXCR5 on TIGIT<sup>-</sup> B cells, and TIGIT<sup>+</sup> B cells were measured by flow cytometry (N = 5 to 10). Representative histograms or dot plots for each marker are shown below the summarized graphs. Each dot represents individual donor and two-tailed unpaired t-test was performed for statistical comparison between TIGIT<sup>-</sup> B cells, and TIGIT<sup>+</sup> B cells. ns: not significant.

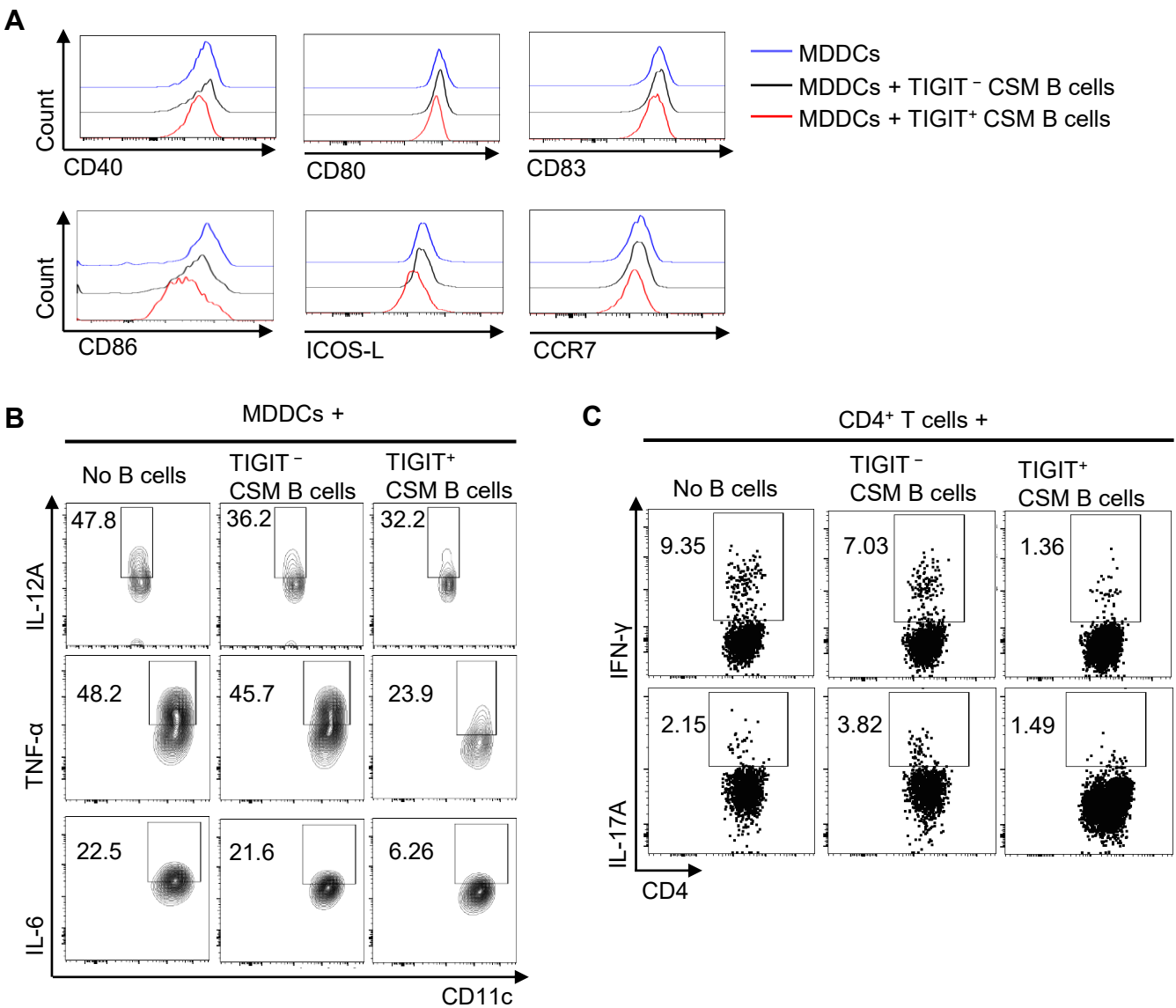

**Figure S8. Representative flow cytometry data associated with Figure 6.**

(A) Representative histograms showing reduced expression of maturation and co-stimulatory markers (CD40, CD80, CD83, CD86, ICOS-L, and CCR7) on LPS-stimulated MDDCs cultured with TIGIT<sup>+</sup> CSM B cells compared with TIGIT<sup>-</sup> CSM B cells or MDDCs alone. (B) Representative FACS plots showing intracellular IL-12A, TNF-α, and IL-6 expression in MDDCs under the same conditions as in A. (C) Representative FACS plots showing intracellular IFN-γ and IL-17A expression by CD4<sup>+</sup> T cells cultured with TIGIT<sup>-</sup> or TIGIT<sup>+</sup> CSM B cells.

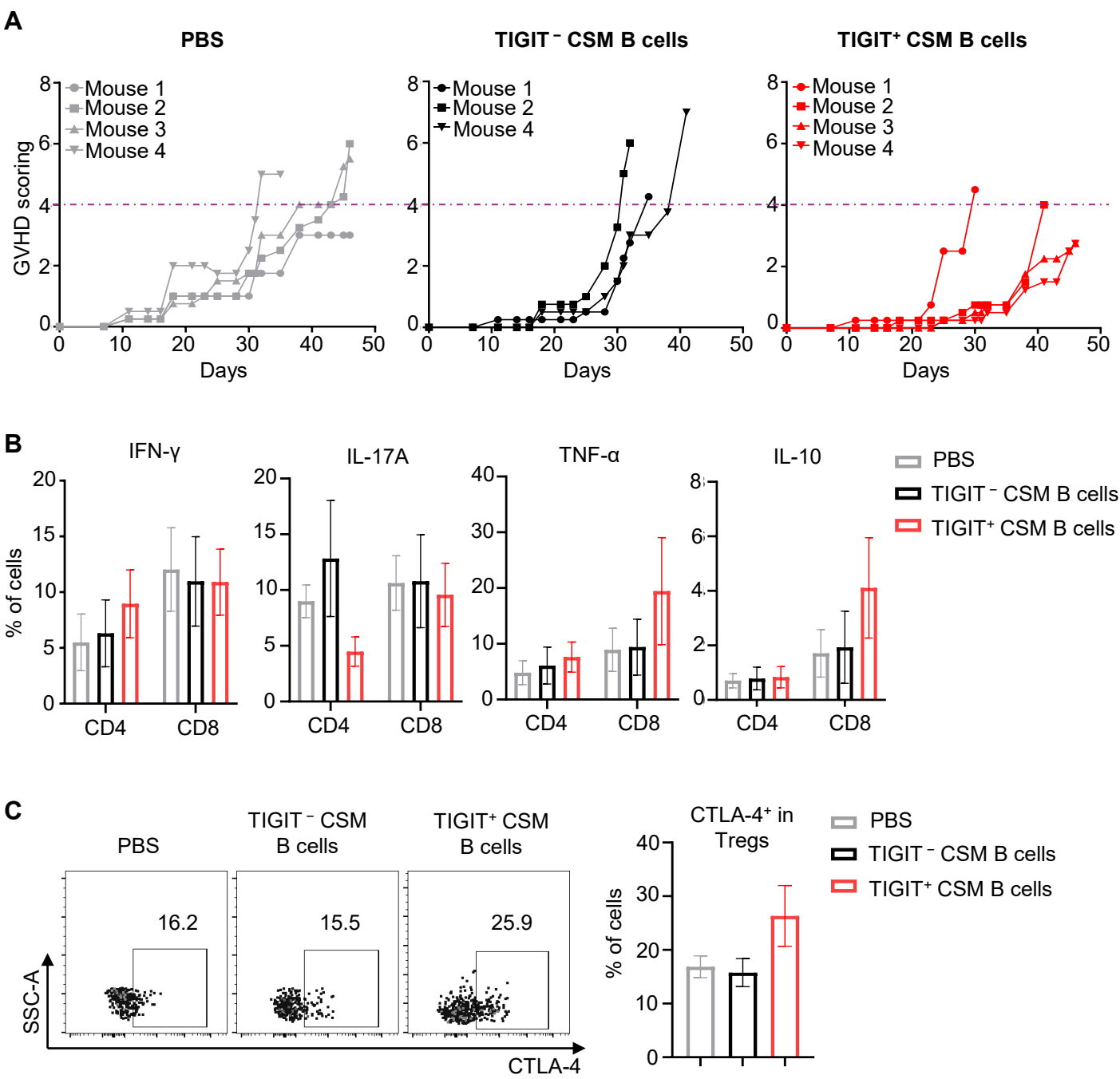

**Figure S9. TIGIT<sup>+</sup> CSM B cells suppress aGVHD progression in NOG mice transplanted with human PBMCs. Data associated with Figure 7.**

(A) GVHD clinical scoring of individual mice from experiment in **Figure 7C-D**. (B) Intracellular IFN-γ, IL-17A, TNF-α, and IL-10 expression in CD4<sup>+</sup> and CD8<sup>+</sup> T cells isolated from spleens. (C) The frequency of surface CTLA-4<sup>+</sup> cells in CD4<sup>+</sup>FOXP3<sup>+</sup>Tregs was measured by flow cytometry.

**Table S1.** Characteristics of human colon tissue samples obtained from patients used in this study.

| Demographics | Patients (n=5)<br>Mean ± SEM or N |
| --- | --- |
| Age (years) | 55 ± 7.64 |
| Gender: |  |
| Female | 3 (60%) |
| Male | 2 (40%) |
| Diagnosis: |  |
| Crohn's disease | 2 |
| Diverticulitis | 2 |
| Obstruction from colonic stricture | 1 |
| Resected Segment: |  |
| Sigmoid Colon | 3 |
| Terminal Ilium | 1 |
| Prior Iliocolic anastomosis | 1 |
